# The spatial landscape of Cancer Hallmarks reveals patterns of tumor ecology

**DOI:** 10.1101/2022.06.18.496114

**Authors:** Mustafa Sibai, Sergi Cervilla, Daniela Grases, Eva Musulen, Rossana Lazcano, Chia-Kuei Mo, Veronica Davalos, Arola Fortian, Adrià Bernat, Margarita Romeo, Collin Tokheim, Enrique Grande, Francisco Real, Jordi Barretina, Alexander J Lazar, Li Ding, Manel Esteller, Matthew H Bailey, Eduard Porta-Pardo

## Abstract

Tumors are complex ecosystems with dozens of interacting cell types. The concept of Cancer Hallmarks distills this complexity into a set of underlying principles that govern tumor growth. Here, we exploit this abstraction to explore the physical distribution of Cancer Hallmarks across 63 primary untreated tumors from 10 cancer types using spatial transcriptomics. We show that Hallmark activity is spatially organized–with 7 out of 13 Hallmarks consistently more active in cancer cells than within the non-cancerous tumor microenvironment (TME). The opposite is true for the remaining six Hallmarks. Additionally, we discovered that genomic distance between tumor subclones correlates with differences in Cancer Hallmark activity, even leading to clone-Hallmark specialization in some cases. Finally, we demonstrate interdependent relationships between Cancer Hallmarks at the junctions of TME and cancer compartments. In conclusion, including the spatial dimension, particularly through the lens of Cancer Hallmarks, can improve our understanding of tumor ecology.

**Significance:** We explored Cancer Hallmarks in 63 primary untreated tumors from 10 cancer types using spatial transcriptomics. This study unveiled spatial patterns in Hallmark activity, with some being more active in cancer cells and others in the non-cancerous tumor environment. Genomic distance impacted Hallmark activity, and we identified interdependencies at the TME-cancer junctions, improving our understanding of tumor ecology.

## Introduction

Over the last few years, we have witnessed an ever-increasing array of discoveries describing the molecular origins of oncogenesis. Among others, we now have catalogues of which genes are driving cancer when mutated^1^, how they interact with the immune system^2^ or new clinically-relevant cancer subtypes based on gene expression profiles^3^. However, the specific molecular changes driving oncogenesis depend on a myriad of variables, such as the specific cancer type^4^, the biological sex of the patient^5^, their germline background^6^, or their age^7^. In the context of this “endless complexity”, Cancer Hallmarks emerge as a powerful integrative approach to rationalize these mechanisms^5^, because they distill the myriad molecular aberrations observed in a tumor into logical principles that describe the necessary features to become malignant^8–10^. For example, while the exact molecular mechanisms might differ from one tumor to the other, driven by different somatic mutations or overexpression of different pathways, all tumors need to “Sustain proliferative signaling”, “Avoid immune destruction” or “Evade growth suppressors’’, just to name three of the various Cancer Hallmarks. In fact, the concept of Cancer Hallmarks is useful to the extent that many of the currently available cancer drugs target specific Hallmarks, such as inhibitors of cyclin-dependent kinases to prevent cancer cells from ‘Sustaining proliferative signaling’ anti-CTLA4 and anti-PD-L1 antibodies to prevent them from ‘Avoiding immune destruction^9, 11^.

Cancer Hallmarks have also been applied to rationalize diverse types of omics datasets, such as those generated by The Cancer Genome Atlas^12^, or to understand the genome-wide effects of individual genes on tumors^1, 13^. The typical process for identifying which Hallmarks are active in certain tumor types or patient samples is by associating them with specific genes, biological pathways, or functional properties^14^. However, until recently, most omics datasets were generated from high-throughput profiling of bulk tumors, limiting our understanding of the interplay between different cell types that form the two major tumor compartments: cancer cells and their surrounding tumor microenvironment (TME). Despite the difficulty in disentangling the cellular origin of the activities of Cancer Hallmarks, the importance of the TME in studying them was already evident when Hanahan and Weinberg, more than a decade ago, expanded their original six Cancer Hallmarks so that, among others, this conceptual framework could be applied to “expand upon the functional roles and contributions made by recruited stromal cells to tumor biology”^9^.

In that sense, the advent of single-cell technologies initiated a new era promising to untangle the complex ecosystem of tumors. The analysis of single-cell transcriptomics from human tumors has revealed an unforeseen richness of different cell types and states in the tumor microenvironment, including different types of immune, stromal, or fibroblast cells^15^. Moreover, such complexity seems to also apply to cancer cells themselves. In most tumors, there seems to be a great variety of cancer cells with multiple transcriptional states, some of which explore, for example, various degrees of differentiation or responses to different stimuli such as hypoxia or interferon^16^. It is worth noting that this transcriptional heterogeneity opens the possibility that Cancer Hallmarks do not need to be assembled by all individual cancer cells, but rather through the cooperation of multiple cell populations^16^.

However, one key piece of information that is lost in single-cell experiments is the actual tissue structure and the context where these different cell types are located. Here is where spatial transcriptomics has provided yet another step forward by allowing us to explore this transcriptional heterogeneity while keeping the spatial distribution of the cells in the original tissue^17^. Among others, the use of spatial transcriptomics has revealed that the area between the tumor and healthy tissue contains a specialized subset of cancer cells^18^, the location of niches of cancer cells with specific functions^19^, or that the tumor edges of renal cell carcinomas are enriched in cancer cells undergoing epithelial-mesenchymal transition^20^.

Here, we quantified how the activity of Cancer Hallmarks in 63 tumor samples from 10 cancer types is distributed through space using spatial transcriptomics. Our findings demonstrate that the distribution of Cancer Hallmark activities is not random, but instead follows a spatial pattern. We also found that cell identity plays a crucial role in determining the activity of these Hallmarks, with some showing consistently higher activity in cancer cells while others being more active in the TME. Our data also indicate that somatic mutations have a limited impact on the variability of Cancer Hallmark activities in most cancers but, given sufficient genomic heterogeneity, some cancer clones can specialize in specific Cancer Hallmarks. Finally, we show that the architecture of Cancer Hallmarks across the entire tumor is largely shaped by the spatial interdependence between phenotypes, as described by the Hallmarks, of cancer cells and those of the TME.

## Results

### Cancer Hallmarks are spatially organized

We have assembled one of the largest Pan-Cancer spatial transcriptomics (VISIUM 10x platform) datasets to date, with 63 primary untreated tumor samples from 10 major cancer types: breast (n=9), prostate (n=9), lung (n=7), brain (n=5), colorectal (n=8), ovary (n=5), bladder (n=5), liver (n=5), pancreas (n=5) and kidney (n=5). Of these, 32 samples were collected from public repositories, while we generated the data for the remaining 31 samples (**Figure 1A, Table S1**) that we ensured were of sufficient high quality (**Figure S1A**, **Table S2**). Overall, our spatial transcriptomics cohort is representative of the most common cancer types.

**Figure 1.**
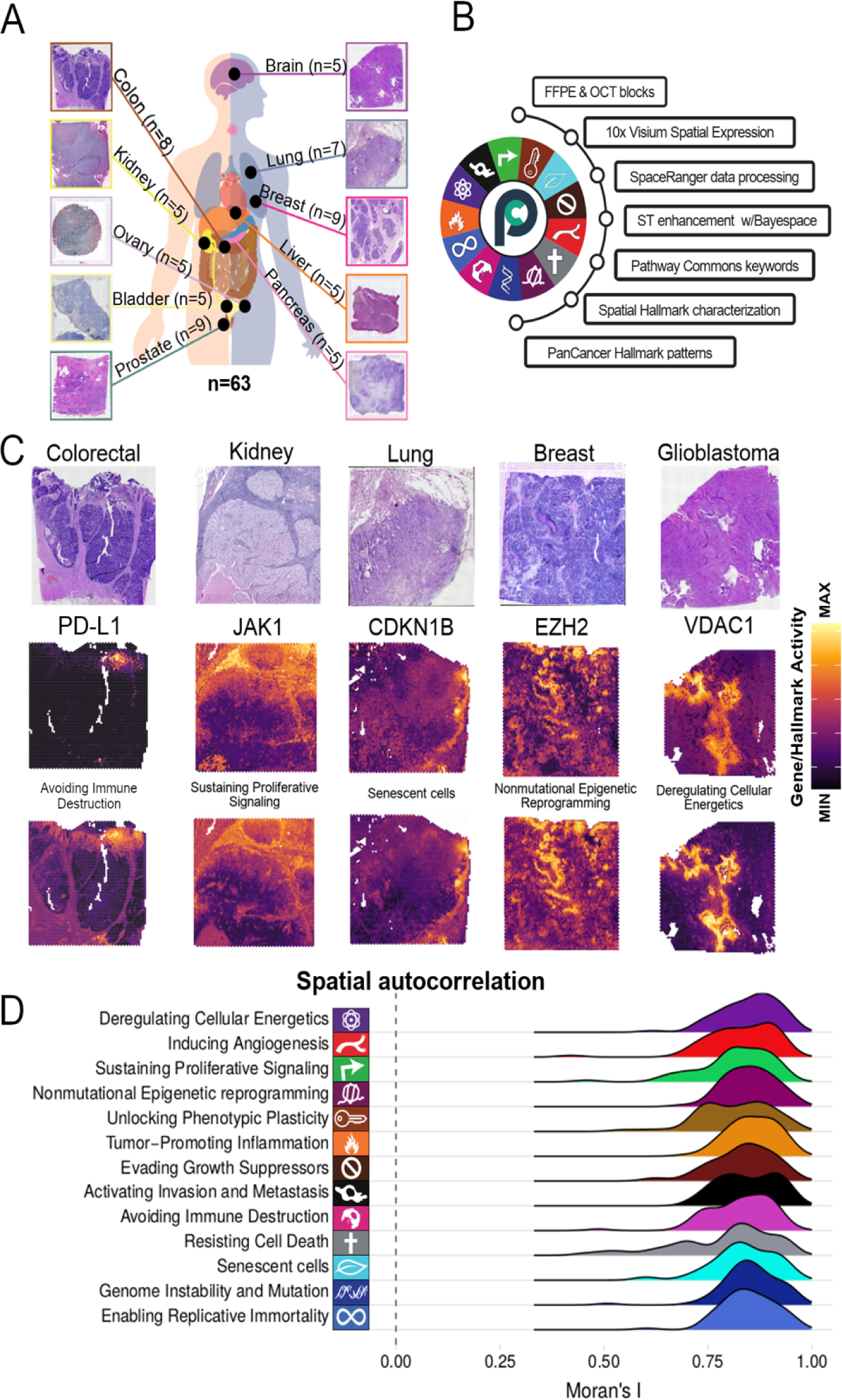
Quantifying the activity of Cancer Hallmarks across space. **A)** Illustration of the spatial transcriptomics cohort highlighting the number of samples for each studied tumor type. **B)** Workflow for b ilding and quantifying each Cancer Hallmark signature in each spatial transcriptomics dataset at near single-cell resolution, and the study of Pan-Cancer patterns. Cancer Hallmark icons are labeled on the outside of the circle. Pathway commons logo is shown at the center of the circle. The article outline is shown to the right of the circle. **C)** A visual comparison among the H&E images (top), the expression of a Hallmark-associated marker (center), and the Hallmark’s activity (bottom) in five of the studied cancer types. **D)** Ridgeplot showing the distribution of Moran’s I across all 63 samples for each of the 13 Cancer Hallmarks. Higher Moran’s I values show non-random clustering of the feature, whereas lower values random distribution of the show eature throughout the space of the tissue.

Recently, the importance of a multi-gene transcriptional program to describe cancer cell states has recently been studied and emphasized^15^. Following this idea, we set out to create transcriptional signatures that describe each of the different Cancer Hallmarks (**Figure 1B**). To that end, we manually curated pathways from the Pathway Commons database to generate gene sets associated with each of the 13 Cancer Hallmarks (all except “Polymorphic Microbiomes”, **Methods**). This ensured that the association of pathways to each Cancer Hallmark was on a conceptual basis, leveraging the collective knowledge of the cancer research community. We established these associations taking into account a trade-off between the considerable biological intertwining Cancer Hallmarks have with one another and their individual specific roles. For example, to quantify the activity of the “(Non)mutational epigenetic reprogramming” Cancer Hallmark we created a gene set that integrates pathways such as “HDACs deacetylate histones”, “Chromatin modifying enzymes” or “HATs acetylate histones”. Individually, each of these pathways only contributes to the Hallmark in a subset of the samples (**Figure S1B**), but when integrated in a single gene expression signature, we can quantify the Hallmark across all samples. The exact pathways and genes included in each Cancer Hallmark are described in **Tables S3 and S4**, respectively.

The number of pathways associated with each Hallmark ranged between 4 and 15 (**Figure S1C**), while the number of genes per Hallmark ranged between 150 and 600 (**Figure S1C**). In total, the signatures for all 13 Hallmarks have 2,699 unique genes, and over 62% of them are associated with a single Hallmark (**Figure S1D**). This balances the required specificity of each gene set while accounting for the biological intertwining across Hallmarks. Finally, we scored the activity of each Hallmark’s gene set in the different samples. Note that, to approach single-cell resolution, we computationally enhanced our spatial transcriptomics data with BayesSpace^21^, which splits each VISIUM spot (which have a ∼55μm diameter) into 6 sub-spots (each ∼20μm in diameter). After scaling and centering these scores within each sample, the final score is what we refer to as the “Hallmark activity” for the rest of the manuscript.

To ensure that the gene lists of each Cancer Hallmark are robust and that their activity can be appropriately quantified throughout all samples, we next calculated the fraction of genes from each Cancer Hallmark signature that were captured in each VISIUM dataset. Our results show that, on average, over 80% of the genes in a Hallmark signature are captured across all samples (**Figure S1E**). Next, we tested the robustness of the Cancer Hallmark signatures to two types of perturbations: missing critical genes (i.e. what happens if we have dropouts of important genes in the signature) and adding potentially redundant genes (i.e. genes that belong to pathways that are very similar to those of each Hallmark and that, therefore, we might have had to include in the original).

For the first control, we correlated the original Cancer Hallmark scores with those of control gene sets that were composed by randomly excluding a number of genes from each Hallmark-associated pathway (**Methods**, **Figure S1F**). The number of excluded genes ranged between 25% and 50% per pathway and, for each fraction that we tested, we repeated the experiment 5 times. For the second control, we correlated the original Hallmark scores with a number of control gene sets that were each composed of the original gene signature plus a number of unique genes from potentially important pathways for the tested Hallmark (**methods**). In both of these control experiments, the correlation coefficients were quite high, with R coefficients ranging between 0.9 and 0.99 (**Figure S1G**). As a final control, we also compared our Hallmark activities with expression patterns of individual genes that would traditionally be associated with specific Cancer Hallmarks (**Figure 1C**). A high-level visual comparison shows how both are positively correlated with each other to various degrees across samples of different cancer types, while also observing nuanced differences across the space of the tissue (**Figure 1C**).

Being confident in our method to quantify the Cancer Hallmark activity, we finally proceeded to their analysis in the spatial transcriptomics dataset. Our first observation was that the activity of all 13 Cancer Hallmarks seems not to be randomly distributed through space but instead restricted to specific areas of the tumor. To quantify this phenomenon, we used a statistic that captures the spatial autocorrelation of any feature: Moran’s I. This statistic ranges between −1 and 1, with features repelling each other in space tending to −1, those with random distribution across space tending to zero, and those with autocorrelation (i.e., tend to form clusters in space) tending to one. The average Moran’s I of all Hallmarks in all samples was around 0.8, confirming that Cancer Hallmarks are spatially clustered and follow a non-random architecture (**Figure 1D, Figure S1H**).

### The activity of Cancer Hallmarks follows a Pan-Cancer tumor architecture

One potential factor contributing to the spatial organization of Cancer Hallmarks is cell identity: cancer cells and the TME can potentially contribute differently to each Hallmark. In fact, while most analyses of the Cancer Hallmarks have focused on cancer cells themselves there has been a growing appreciation for the potential contribution to some Hallmarks by cells from the TME^9^. To explore this in detail, we classified each subspot in each tumor sample into one of the two major tumor compartments: cancer cells and the TME with ESTIMATE^22^ (**Methods**). This tool uses a gene expression signature to quantify the presence of cancer cells in an RNAseq dataset. We clustered the ESTIMATE scores of each sample into five groups, where spots from Cluster 1 have low ESTIMATE scores (suggesting a high content of cancer cells), and spots that belong to Cluster 5 have high ESTIMATE scores (suggesting a low content of cancer cells, i.e. mostly TME, **Figure 2A**). We validated the accuracy of these assignments by quantifying the presence of cancer and stromal cells in each cluster with histopathological annotations from certified pathologists that used good-quality Hematoxylin and Eosin (H&E) images for 17 samples from 5 cancer types (**Figure 2B, Figure S2A, Table S5**). The results show an excellent agreement between ESTIMATE-based clusters and the percent of cancer cells annotated by human pathologists, with spots in Cluster 1 having, on average, >90% of cancer cells and <5% of stromal cells, and spots in Cluster 5 having, on average, <10% of cancer cells and >85% of stromal cells (**Figure 2B**).

**Figure 2.**
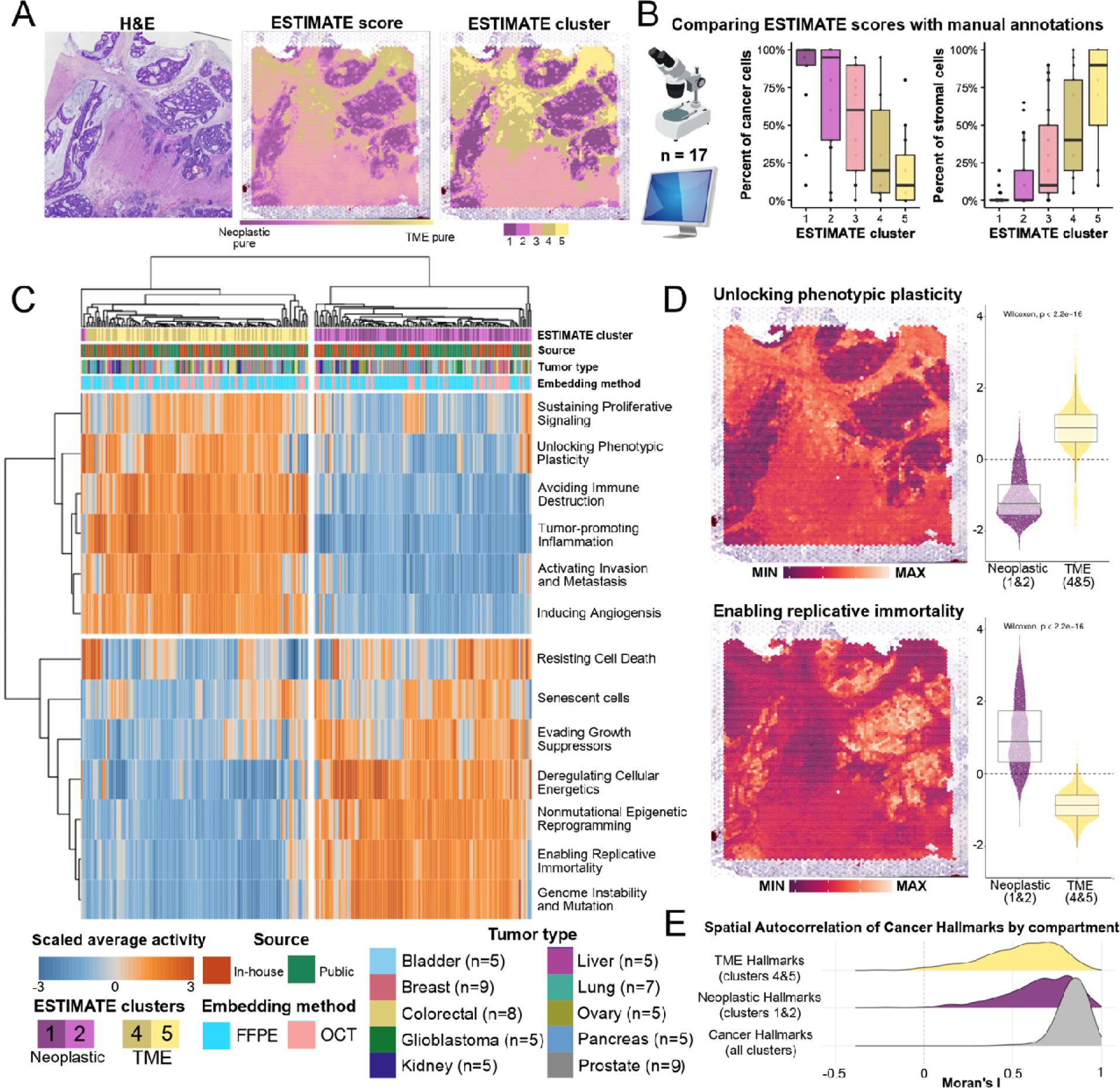
The Pan-Cancer architecture of Cancer Hallmarks. **A**) H&E of a colorectal cancer sample (left) with the computed ESTIMATE scores for each individual subspot (center), ranging from pure neoplastic cell content in purple to pure TME content in yellow. The score is categorized into five clusters by applying k-means (right). **B**) Benchmark of ESTIMATE scores on 17 samples from 5 cancer types annotated by human pathologists to quantify the percentage of stromal and cancer cells within each cluster. **C**) A 2D-hierarchically clustered heatmap showing the scaled average Cancer Hallmark scores (rows) within a cluster in each sample (columns). The annotation layer shows from top to bottom: (1) the ESTIMATE cluster with 1 and 2 corresponding to the neoplastic compartment, while 4 and 5 corresponding to the TME compartment, (2) the source of the sample whether it was publicly available or in-house generated, (3) the tumor type of the sample, and (4) the tissue embedding method. **D**) Distribution of Hallmark scores in neoplastic (clusters 1&2) and TME (clusters 4&5) compartments as well as their spatial patterns in the previously shown colorectal sample for “Unlocking phenotypic plasticity” (top) and “Enabling replicative immortality” (bottom) Hallmarks. **E**) Distribution of Moran’s I computed for all Hallmarks across the whole tissue of each sample, for the 7 Hallmarks whose activities are differentially higher in the neoplastic compartments (in clusters 1&2), and for the 6 others whose activities are differentially higher in the TME compartments (in clusters 4&5).

We then compared the average Cancer Hallmark activity of the spots from each of the different ESTIMATE clusters across all samples from all tumor types. To ensure that we are comparing spots containing mostly cancer cells or TME, we excluded spots from Cluster 3 from downstream analyses as it includes all spots with intermediate ESTIMATE scores of each sample, which can potentially be a “buffer-like” zone that containing intermixed proportions of cancer cells and cells from TME depending on the sample. Our results show a clear Pan-Cancer architecture, with Hallmark activities clustering by which tumor compartments are more active (p-value < 0.0001, Fisher’s Exact Test). Notably, these clusters did not group by cancer type (p-value = 1, Fisher’s Exact Test), tissue embedding method (p-value = 1, Fisher’s Exact Test), or source of the data (p-value = 0.9, Fisher’s Exact Test). These results show that cell identity plays a key role in shaping the spatial architecture of Cancer Hallmarks.

We found 7 Hallmarks more active in ESTIMATE clusters 1 and 2 (i.e., cancer cells, **Figure 2C**). These cancer-cell associated Hallmarks are: “Evading Growth Suppressors’’, “Enabling Replicative Immortality”, “Deregulating Cellular Energetics’’, “Senescent cells”, “(Nonmutational) Epigenetic Reprogramming’’, “Genomic Instability and Mutations’’, and “Resisting Cell Death”. In some exceptional cases, activities of “Senescence” or “Resisting Cell Death” could also be active enough in TME compartments of some samples, suggesting the presence of senescent immune or stromal cells with induced survival mechanisms that may contribute to the malignant phenotype^10^.

The 6 other Hallmarks were more active in spots from ESTIMATE Clusters 4 and 5, which correspond to the TME (**Figure 2C**). Among others, these include “Inducing Angiogenesis”, “Activating Invasion and Metastasis”, “Unlocking Phenotypic Plasticity”, whose putative functional roles are to modulate the extracellular matrix and neighboring blood vessels to facilitate the extravasation of cancer cells by acquiring stemness properties through processes such as epithelial-to-mesenchymal (EMT) transition. The TME activities of another two Hallmarks - “Avoiding Immune Destruction’’ and “Tumor-Promoting Inflammation”- indicate the participation of immune cells, as well as secreted signaling molecules by cancer cells such as the senescence-associated secretory phenotype (SASP) into the surroundings, all of which concurring the activity of these two Hallmarks being mainly part of the TME. Finally, “Sustaining Proliferative Signaling” is also more active in the TME as the cancer cells continuously need extrinsic signals such as mitogens from the TME to sustain the proliferative capacity of cancer cells.

Notably, even if the activity of all Cancer Hallmarks showed marked differences between compartments (**Figure S2B**), there is still large variability in the activity of spots within the same compartment. For example, the activity of “Unlocking Phenotypic Plasticity” in the TME compartment from one colorectal cancer sample showed high activity in 25% of TME-spots. Notably these spots were not randomly dispersed throughout the TME, but rather maintained spatial order (**Figure 2D, top**). Similarly, using the same colorectal sample, “Enabling Replicative Immortality” was extraordinarily high in 25% of the neoplastic-spot, and these spots also spatial cohesive (**Figure 2D, bottom**). Specifically, the activity of these two Hallmarks in each compartment was spatially restricted for neoplastic and TME compartments (Moran’s I = 0.61 and = 0.66, respectively). A systematic exploration of the spatial distribution of all Hallmarks within each compartment revealed that the activity of the Hallmarks is not randomly distributed within the TME and cancer compartments: the average Moran’s I of the 7 neoplastic Hallmarks within the neoplastic compartments and that of the 6 TME Hallmarks within the TME compartments across all 63 samples were 0.64 and 0.55, respectively (**Figure 2E, Figure S2C**). Also, the Moran’s I values for each Hallmark were also significantly higher than those of random gene signatures even for “Resisting Cell Death” (**Figure S2D**). This suggests that, while cell identity plays a major role in determining the overall activity of Cancer Hallmarks and their spatial distribution within tumors, other factors must explain the internal spatial distribution of Cancer Hallmarks within the neoplastic and TME compartments.

### The interplay between genetic and phenotypic variability throughout space

Another factor that could be contributing to the spatial organization of Cancer Hallmarks’ activities is the presence of different cancer clones. During their evolution, cancer cells can acquire different somatic mutations, which might, in turn, lead to transcriptional and phenotypic differences between clones, including differences in the activity of Cancer Hallmarks^23^. To explore this hypothesis, we used inferCNV (**Methods**) to find tumor areas with different CNV states (inferred from their gene expression profiles) as evidence that they could be different genomic clones (**Figure S3**)^24^. Overall, InferCNV detected a total of 693 clones across all samples. Next, we calculated the genetic distance of each pair of clones in each sample. We did so based on the fraction of the genome that differs in CNV state between both clones. Finally, we calculated the correlation between the genetic distance of all 2740 pairs of clones and the difference in the activity of different Cancer Hallmarks (**Figure 3A**).

**Figure 3.**
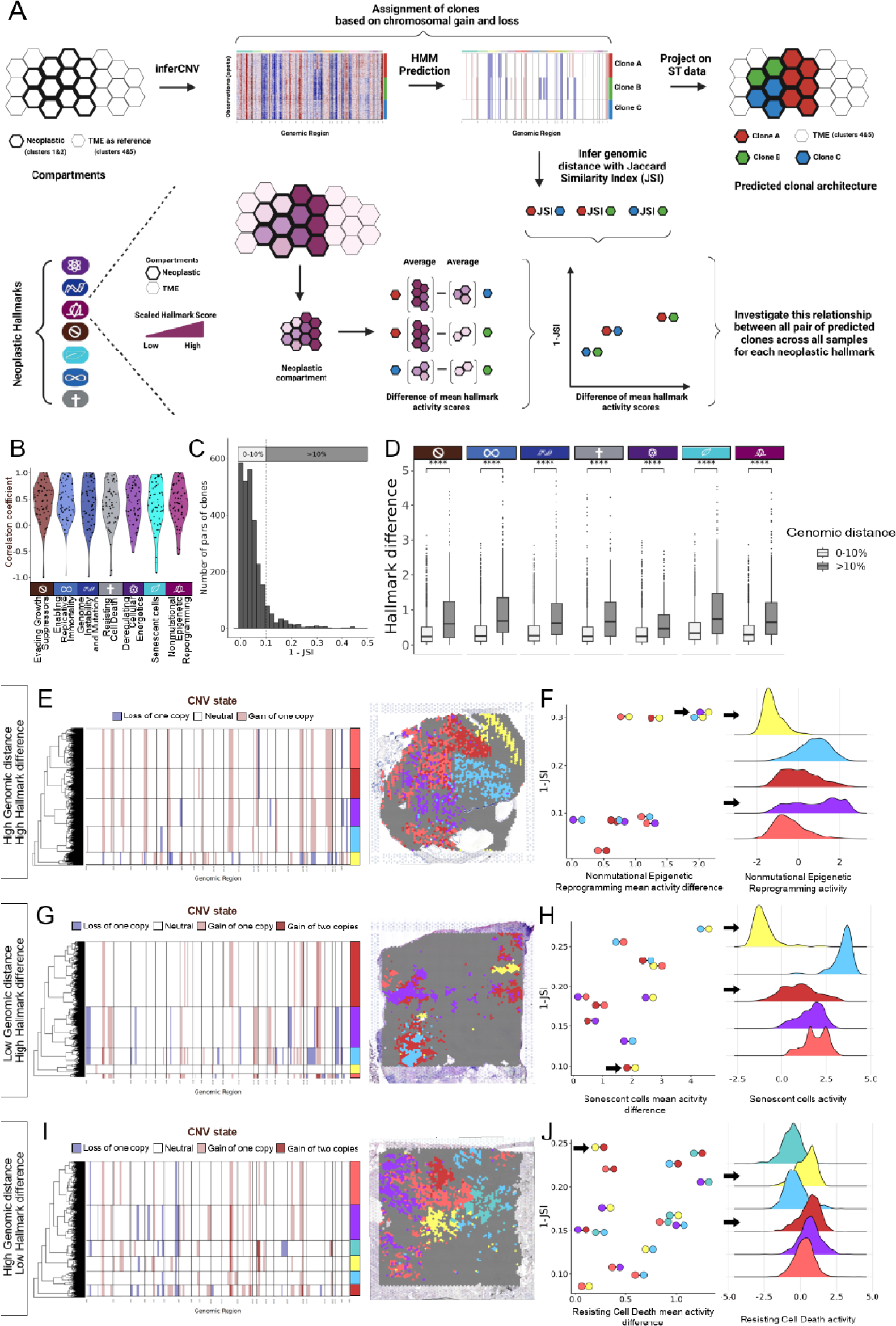
Genomic variation explains the variability of Cancer Hallmark activities to some degree. **A)** A schematic diagram illustrating the process of inferring CNV status of reported clones predicted by inferCNV on a given spatial transcriptomics sample, and calculating the genomic distance between all pairs of clones to be compared to the difference in the activity of each neoplastic Hallmark between the corresponding pair of clones across all samples. **B)** Violin plots of Pearson correlation coefficients as a result of correlating the genomic distances (measured by 1-Jaccard Similarity Index) and the difference in each neoplastic Hallmark activity between all pairs of clones across all samples. **C)** A histogram of the distribution of the genomic distances between all pairs of clones across all samples categorized in two bins: those having 0-10% >10% genomic distance. **D)** Boxplots comparing the difference in activities of neoplastic Hallmarks between both categories of genomic distances. HMM predictions of genomic states reported by inferCNV and the projected clones on the histology images in an ovarian cancer sample (**E**), a Pancreas cancer sample (**G**), and a lung cancer sample (**I**). Each of these samples exemplify one of three scenarios: **F)** a pair of clones colored in purple and yellow from the sample shown in **E** exhibit high genomic distance and high difference in the activity of “Nonmutational Epigenetic Reprogramming” **H)** a pair of clones colored in red and yellow from the sample shown in **G** exhibit low genomic distance and high difference in the activity of “Senescent cells” **J)** a pair of clones colored in red and yellow from the sample shown in **I** exhibit high genomic distance and low difference in the activity of “Resisting Cell Death”. ns: p > 0.05, *: p <= 0.05, **: p <= 0.01, ***: p <= 0.001, ****: p <= 0.0001

Our results show that there is an overall positive correlation between how different two clones are at the genetic level, and how much they differ in their Hallmark activities (Pearson correlation coefficients ranging from 0.3 to 0.4 on average, **Figure 3B**). Furthermore, the skewed distribution of genomic distances between all pairs of clones across all samples suggests that the vast majority of these pairs have a relatively small genomic distance (**Figure 3C**). We categorized this distribution of genomic distances into two bins: 0-10% and > 10% of difference in the genome (**Figure 3C**). Consistent with the previous observation, the differences in the mean activity of each of the 7 neoplastic Hallmarks were significantly higher (p values < 0.05) in pairs of clones with higher genomic distance compared to those with lower genomic distance (**Figure 3D**). When focusing on individual Hallmarks, the overall differences in the mean activity of “Deregulating Cellular Energetics” between pairs of clones is generally lower (< 1 standard deviation on average) compared to that of other neoplastic Hallmarks. This suggests that cancer cells need a similar activity of this Hallmark regardless of their genetic alterations, making this Hallmark, potentially, a requirement for all cancer cells.

Our results reveal four possible scenarios at the sample level depending on the differences at the genomic and Hallmark levels. The first and most common one is the “low-low” scenario, where two clones with low genomic distance also have low differences in their Hallmark activities. We observed this in 2261 pairs of clones across all samples (82.52% of the total pairs), and in at least one pair of clones in 92% of samples. The second scenario is that of high genomic distance and high difference in Hallmark activity (i.e. “high-high”). In this scenario, the acquisition of genetic alterations leads to the specialization of certain clones in specific Hallmarks. We found one such pair in one of the ovarian cancer samples **(Figure 3E**). In this sample inferCNV predicted 5 distinct clones based on the HMM model. The genomic distances between each pair of the 5 clones varied from low (≤10%) to high (≥30%) (**Figure 3F**). The group of clones with low genomic differences also exhibited similar activities in the “Nonmutational Epigenetic Reprogramming” Hallmark, while the latter exhibited medium to high differences in the activity of the same Hallmark. Notably, the yellow and purple clones are the most divergent in terms of their genomic distance and the Hallmark difference (**Figure 3F**). Overall, this pattern was rare across pairs of clones, as we only observed it in 93 pairs across all samples (3.39% of the total pairs). However, at the sample level it was relatively common, as these 93 pairs of clones were spread across 40% of all samples.

The other two scenarios include the cases where the difference between a pair of clones in one of the variables is low, but in the other variable is high (high-low and low-high). In this case, it is likely that genomics play a less important role than the interplay between cancer cells and the TME. For example, the third scenario includes pairs of clones that have differences in their Hallmark activity, despite the fact that they are genomically similar (≤10% of genomic distance), which happened in 188 pairs of clones across all samples (high-low, 6.86% of the total pairs), and in at least one pair of clones in 49% of samples in this scenario. This is the case for instance for a pancreas cancer sample (**Figure 3G**), where inferCNV predicted 5 distinct clones based on the HMM model. While the genomic distance between clone pairs in this sample varied from 10% to over 25%, the yellow and red clones were genomically quite similar despite them both exhibiting a high difference (2 standard deviations) in the mean activity of “Senescent cells” Hallmark (**Figure 3H**).

Finally, some clones have similar Hallmark activities despite having high (≥30%) genomic distance (low-high). This scenario suggests phenotypic convergence, which we observed in 198 pairs of clones across all samples (7.23% of the total pairs), and in at least one pair of clones in 49% of samples. These included one of the lung cancer samples (**Figure 3I**), where inferCNV predicted 6 distinct clones based on the HMM model. Notably, the yellow and red clones exhibit very similar activity of “Resisting Cell Death” Hallmark even if they both are genomically distant by about 25% (**Figure 3J**). These two latter scenarios suggest that, while genomic evolution may play a role in shaping the spatial architecture of Cancer Hallmarks, there is likely a significant involvement of cell-extrinsic events that influence the spatial landscape of the neoplastic Hallmarks, potentially through juxtacrine or paracrine signaling from the TME^25^.

### Quantifying the spatial interdependence of Cancer Hallmarks

Another factor that might contribute to the spatial architecture of Cancer Hallmarks is their spatial interdependence: it is possible that the activity of certain Hallmarks is influenced by the location of other Hallmarks. Notably, if such spatial interdependence exists, one of its likely contributing mechanisms is cellular crosstalk. After all, there is ample evidence showing that cancer cells send molecular signals to their surrounding TME and vice versa. However, thanks to the fact that we are using the phenotypic concept of Cancer Hallmarks, we can go one step further and explore such spatial interdependence through the lens of ecological dynamics^26^, where the location of cells with specific phenotypes is shaped by the presence or absence of cells with other phenotypes. This has been recently shown, among others, in colorectal cancer, where cancer cells exhibit different transcriptional programs (and phenotypes) depending on their position relative to the invasive front of the tumor^27^.

To explore such spatial interdependence through the lens of Cancer Hallmarks, we used random forest regression to model how the spatial distribution of the TME Hallmarks predicted the spatial distribution of the neoplastic Hallmarks (**Figure 4A**) and vice-versa (**Figure S4**). To account for the geometrical distribution of the activity of Cancer Hallmarks in either experiment, we translated the activity of the Cancer Hallmarks from the predicting compartment (or predictor) into radar scores within the predicted compartment (or target) (**Methods**). For example, suppose we are trying to model how the Hallmark activities from the TME predict the activity of the Hallmarks from cancer cells. In that case, we transform the activity of the TME into radar scores within cancer cells (**Figure 4A**). Once we have calculated all necessary radar scores for all Hallmarks, we train the model on 80% of randomly selected spots, followed by predicting the target Hallmark activity (response) using the remaining 20% of the spots (**Figure 4A**). The overall workflow to transform the activity of one compartment into radar scores within the other compartment is illustrated in **Figure 4B** for “Avoiding immune destruction” whose TME activity is translated into radar scores within the neoplastic compartment in a kidney cancer sample where the target Hallmark is “Genome Instability and Mutation” in the neoplastic compartment.

**Figure 4.**
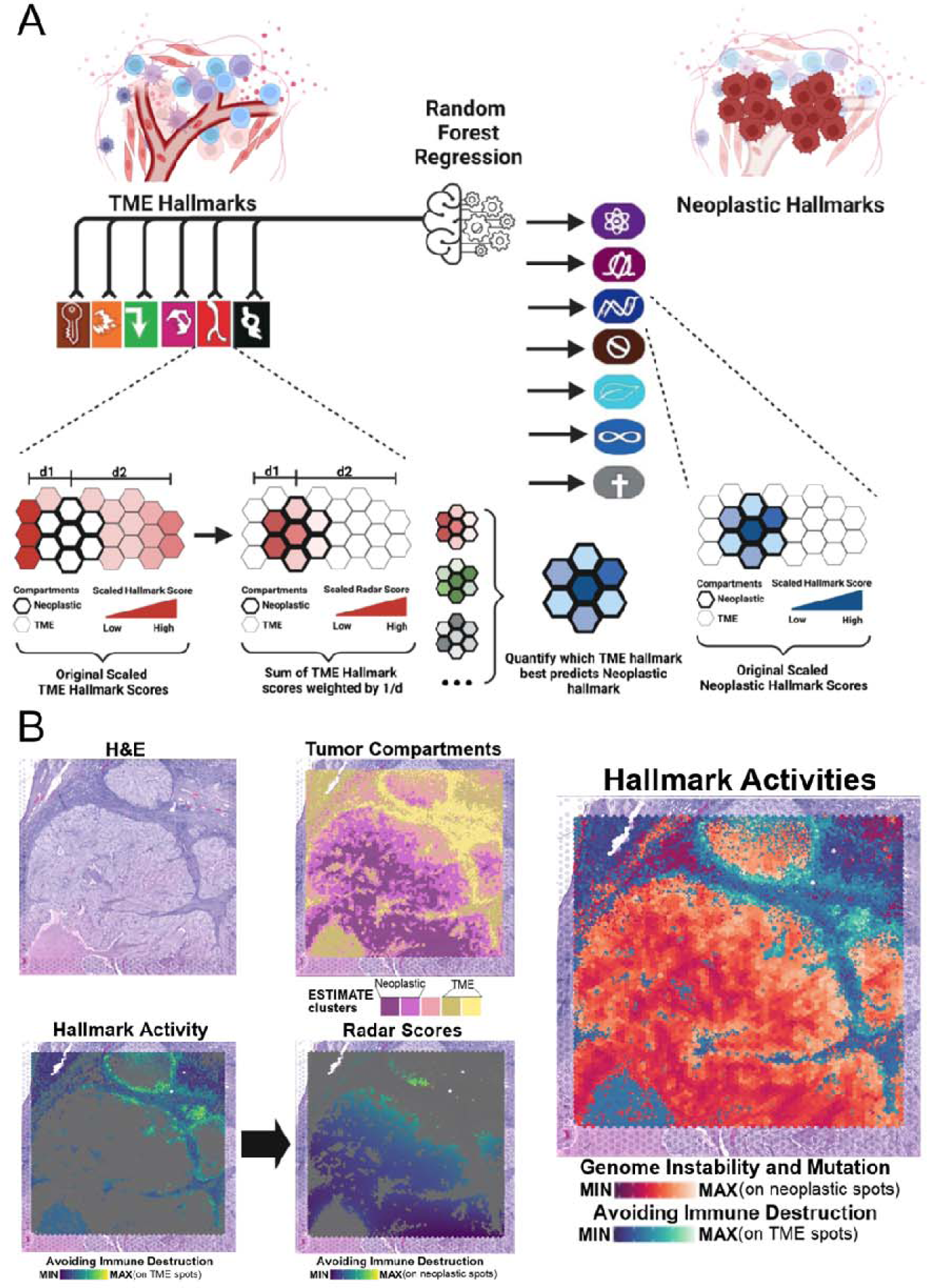
Describing the spatial interdependence between neoplastic and TME Hallmarks. **A**) Illustration of using Random Forest regression to predict each neoplastic Hallmark within the neoplastic compartment using the combination of the TME Hallmarks from the TME compartment through their conversion into “radar” scores within the neoplastic compartment where the targets are located. **B)** Highlighting an example of a model from a kidney tumor in which “Genome Instability and Mutation” is the target and “Avoiding Immune Destruction” is one of the predictors. Top to bottom are depictions of 1) upper row: the tissue’s H&E image (left), and the ESTIMATE clusters (right), 2) bottom row: the activity of “Avoiding immune destruction” on the TME subspots (clusters 4&5) on the left. and the radar score of “Avoiding Immune Destruction” in the neoplastic compartment (clusters 1&2) on the right. The most right panel depicts the activities of “Avoiding Immune Destruction” in TME compartment (in blue) and “Genome Instability and Mutation” in the neoplastic compartment (in red) together in the same ST histology image.

We used the Shapley Additive Explanations (SHAP) values to perform the predictions, which allowed us to identify how the model prioritizes the relationships between the features (radars) and the target Hallmark with two metrics: feature importance and spatial dependency (**Methods**). Feature importance allowed us to rank the predictor Hallmarks of one compartment based on how much they explain the spatial distribution of a target Hallmark activity in the other compartment. Spatial dependency, on the other hand, indicates in what direction a Hallmark is predicting its target. Specifically, a positive spatial dependency implies that both the predictor and its target in their respective compartments tend to be active in proximity throughout the majority of their interface. On the other hand, a negative spatial dependency implies that the activities of both the predictor and its target tend to be far away. Finally, a non-linear spatial dependency may suggest the tendency for the predictor and its target to have similar activities in proximal or distal specific regions throughout the tumor section.

### The tumor microenvironment shapes the spatial architecture of cancer cells through Hallmark collaboration

We first modeled how the activity of Hallmarks from the TME predicted the spatial distribution of Hallmarks within the neoplastic compartment, the results of which are summarized in **Figure 5**. Overall, our random forest models show that the TME Hallmarks have high predictive accuracy (mean R^2^ of 0.71 across all models) of the 7 neoplastic Hallmarks when modeled as the targets (**Figure 5A**). The addition of the relative importance of each predictor across all samples revealed that the predictors (the 6 radars for the TME Hallmarks) of neoplastic Hallmarks showed low variability in their prediction importance, except for “Inducing Angiogenesis” which consistently appeared to be the least important predictor of each target (**Figure 5B, Figure S5A**). This suggests that, in most cases, the spatial distribution of the neoplastic Hallmarks is the result of a combination of multiple factors, including the relative activity of multiple TME Hallmarks.

**Figure 5.**
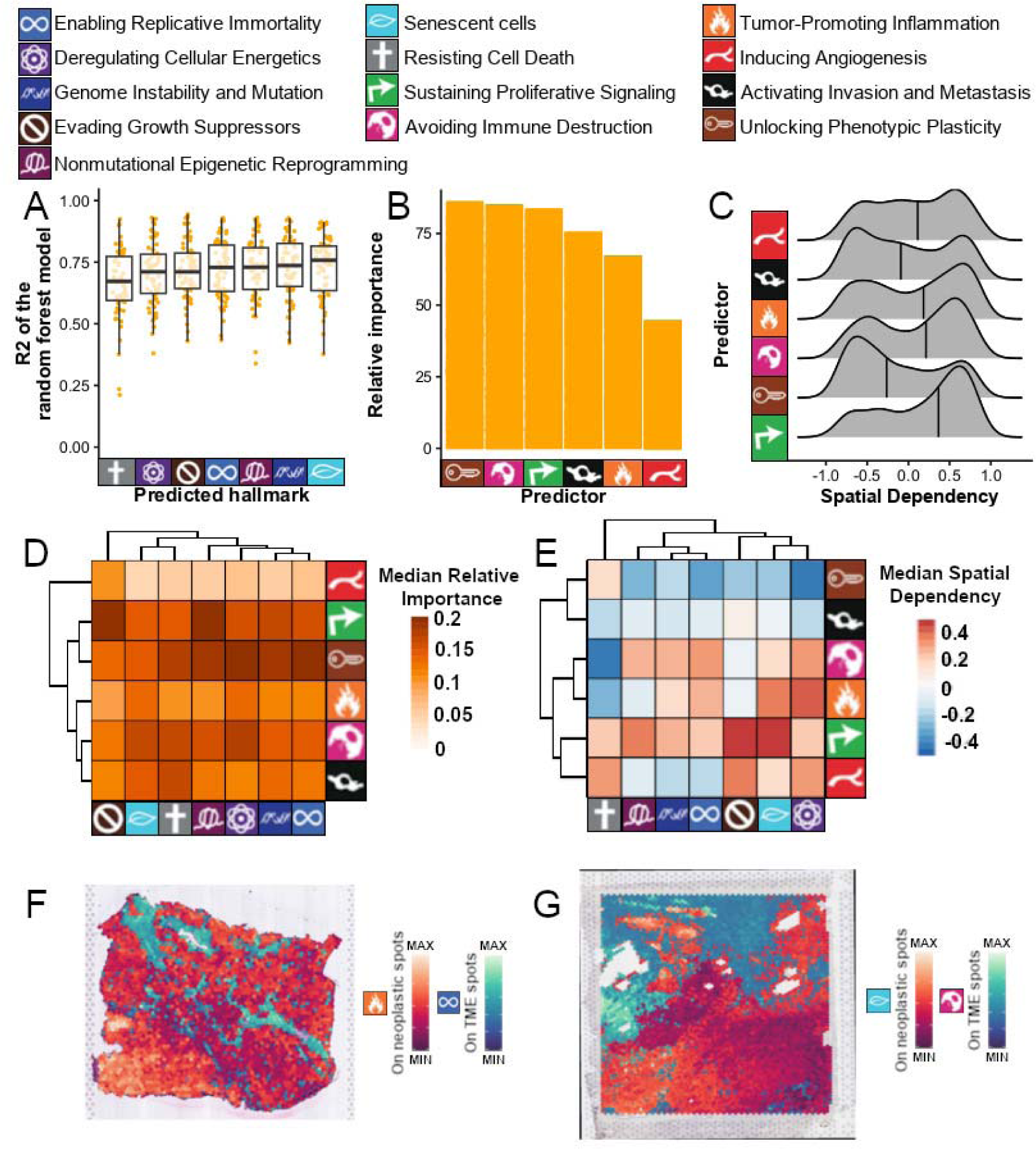
The TME Hallmarks predict the distribution of neoplastic activity through a variety of mechanisms. **A)** Boxplot showing the R2 of each random forest model (y-axis) clustered by the predicted hallmark (x-axis). **B)** Barplot showing the sum of relative importances (y-axis) of each predicting cancer hallmark (x-axis). **C)** Ridge plots of the Pearson correlation coefficients for the spatial dependency (calculated by correlating the predictor and its SHAP values as a proxy to its target), where positive coefficients indicate a positive spatial dependency and negative coefficients indicate a negative spatial dependency. **D)** Heatmap clustering the predictor Hallmarks (rows) according to their relative importance when predicting the target Hallmarks (columns). **E)** Same as **D** but clustering according to the spatial dependency of the predictive model. **F)** The scenario of having high relative importance and positive spatial dependency is illustrated in a prostate cancer sample (with “Avoiding Immune Destruction” being the predictor and “Senescent cells” being the target). **G)** A scenario of having high relative importance and negative spatial dependency is illustrated in a liver cancer sample (with “Tumor-Promoting Inflammation” being the predictor and “Enabling Replicative Immortality” being the target).

Such collaborative influence may be attributed to the heterogeneous cellular make-up of the TME itself. To explore this hypothesis, we quantified the presence of over 20 different cell types in the subspots from the TME compartment (ESTIMATE clusters 4&5)(**methods, Figure S5B**). Next, we calculated the correlation between the activity of the 6 TME Hallmarks with the quantified signatures of each cell type in each spot, allowing us to identify which cell types are contributing to each Hallmark. Among others, this revealed that, as expected, the activity of “Avoiding Immune Destruction” is highly correlated with cells of only the immune system (including T cells, T helper cells, B cells, and monocytes) The activity of the other five Hallmarks, however, showed some unexpected patterns. For example, the activity of “Tumor-promoting inflammation” correlated with cells of the immune system, but also with endothelial cells, fibroblasts, and pericytes. Similarly, the activity of “Activating Invasion and Metastasis” and “Unlocking phenotypic plasticity” correlated with both fibroblasts and pericytes. The activity of “Inducing angiogenesis” also correlated with some expected cell types (such as endothelial cells), but also with cells from the immune system (megakaryocytes, monocytes, etc.), pericytes, and fibroblasts. Finally, the activity of “Sustaining Proliferative Signaling” showed the lowest mean correlation coefficients with individual cell types, suggesting that its activity is spread throughout multiple cell types of all systems. Taken together, these results suggest that many of the TME Hallmarks may not necessarily require cell types from a specific system in order to exhibit their functional dynamics within the tumor, but its rather their collective action that finally determines the overall activity of the Hallmark in the TME, highlighting the importance of measuring phenotypes (such as Hallmarks), instead of individual cell types.

Regarding the spatial dependency, the TME Hallmarks seem to be more variable. For example, “Sustaining Proliferative Signaling” clearly exhibited a tendency to predict its targets by its proximity (as evidenced by its overall high positive spatial dependency). By contrast, “Unlocking Phenotypic Plasticity” showed the opposite tendency by predicting its targets via its distance-that is a high negative spatial dependency with its targets (**Figure 5C**). The other predictors (“Avoiding Immune Destruction’’, “Tumor-Promoting Inflammation”, “Activating Invasion and Metastasis”, and “Inducing Angiogenesis”) do not have a clear pattern and seem to have a tendency to predict their targets by both proximity and distance (**Figure 5C**).

Looking at the detailed relationships between pairs of Hallmarks for both, relative importance (**Figure 5D**) and spatial dependency (**Figure 5E**) revealed further insights. For example, while it is true that “Inducing angiogenesis” is the TME Hallmark with the lowest relative importance, it has a significant contribution when predicting “Evading growth suppressors”, with a strong positive spatial dependency. Another interesting pattern comes from “Avoiding immune destruction”, which has a positive spatial dependency with all neoplastic Hallmarks with the exception of “Resisting Cell Death”, where the spatial dependency is negative. This is consistent, among others, with cancer cells not needing to resist cell death when in contact with an inactive immune response.

To depict the relationship of a TME Hallmark predicting a target neoplastic Hallmark visually, we overlaid the activities of a predictor-target pair on histology images from different cancer types. The case where a TME Hallmark such as “Avoiding Immune Destruction” has a high relative importance in predicting a target neoplastic Hallmark such as “Senescent cells’’ through a positive spatial dependency is illustrated on a Prostate cancer sample (**Figure 5F**). By contrast, an example of a TME Hallmark such as “Tumor-Promoting Inflammation” having a high relative importance in predicting a target neoplastic Hallmark such as “Enabling Replicative Immortality” through a negative spatial dependency is illustrated on a liver cancer sample (**Figure 5G**).

Finally, we inspected the results of the random forest models for specific tumor types. Unlike the PanCancer pattern, which showed similar importance for 5 of the 6 TME Hallmarks, here the TME Hallmarks varied more in their importance at predicting their targets (**Figure S5C**). For example, “Activating Invasion and Metastasis” is the top predictor for neoplastic Hallmarks in bladder cancer and glioblastoma, while “Sustaining Proliferative Signaling” is one of the top predictors in ovarian, breast, and colorectal cancers. Next, we inspected the results of specific tumor types. Unlike the PanCancer pattern, which showed similar importance for 5 of the 6 TME Hallmarks, here the TME Hallmarks varied more in their importance at predicting their targets (**Figure S5C**). For example, “Activating Invasion and Metastasis” is the top predictor for neoplastic Hallmarks in bladder cancer and glioblastoma, while “Sustaining Proliferative Signaling” is one of the top predictors in ovarian, breast, and colorectal cancers.

### Cancer cells configure the spatial architecture of the microenvironment through Hallmark specialization

Next, we modeled how the activity of Hallmarks in the neoplastic compartment can predict the spatial distribution of the Hallmarks in the TME, the results of which are summarized in **Figure 6**. Again, our random forest models showed high predictive accuracy (mean R^2^ of 0.67 across all models, **Figure 6A**). However, unlike the previous experiment, the predictors (the 7 radars for the neoplastic Hallmarks) showed higher variability in their overall prediction importance (**Figure 6B, Figure S6A**). Here there is a clear gradient showing that “Resisting Cell Death” is the most important neoplastic Hallmark when predicting the spatial distribution of the TME. This is followed, in decreasing order of overall importance, by “Deregulating Cellular Energetics”, “Evading Growth Suppressors”, and “Senescent cells”. The last three neoplastic Hallmarks were the least important in predicting their targets were “Genomic Instability and Mutations”, “Enabling Replicative Immortality”, and “(Nonmutational) Epigenetic Reprogramming’’. Furthermore, these predictors also varied in terms of their spatial dependency with their targets. While “Evading Growth Suppressors’’ showed a general tendency to predict its targets in close proximity, “Genome Instability and Mutations” and “Enabling Replicative Immortality” tended to predict their targets with a negative spatial dependency (i.e. when the activity of the neoplastic Hallmark is low, the activity of the TME Hallmark tends to be high, **Figure 6C**). The other four neoplastic Hallmarks do not seem to have any specific pattern, predicting the activity of the TME Hallmarks sometimes through proximity, other times through distance (**Figure 6C**).

**Figure 6.**
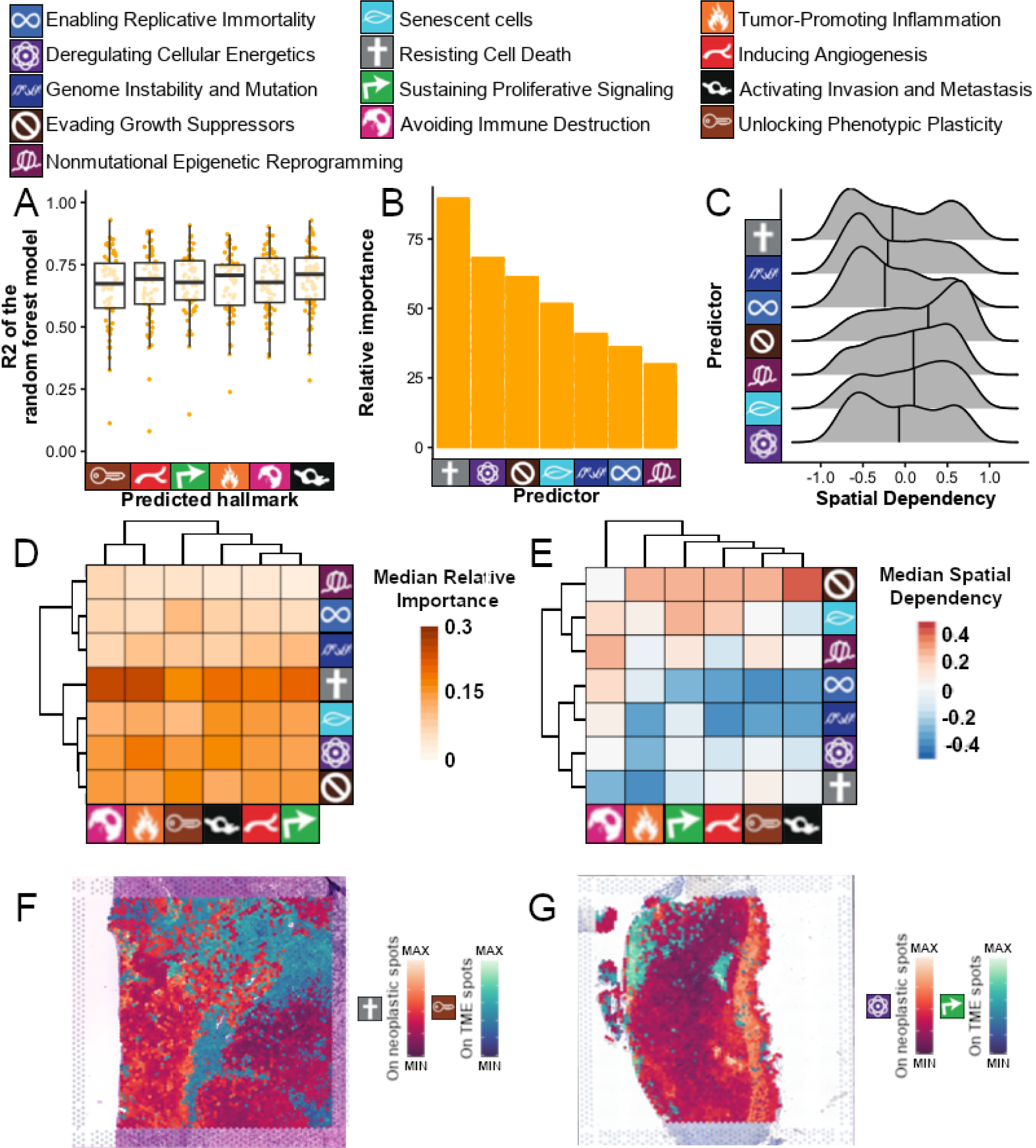
Neoplastic Hallmarks predict the spatial distribution of the TME through specific activities. **A)** Boxplot showing the R2 of each random forest model (y-axis) clustered by the predicted hallmark (x-axis). **B)** Barplot showing the sum of relative importances (y-axis) of each predicting cancer hallmark (x-axis). **C)** Ridge plots of the Pearson correlation coefficients for the spatial dependency (calculated by correlating the predictor and its SHAP values as a proxy to its target), where positive coefficients indicate a positive spatial dependency and negative coefficients indicate a negative spatial dependency. **D)** Heatmap clustering the predictor Hallmarks (rows) according to their relative importance when predicting the target Hallmarks (columns). **E)** Same as **D** but clustering according to the spatial dependency of the predictive model. **F)** The scenario of having high relative importance and positive spatial dependency is illustrated in a kidney cancer sample (with “Resisting Cell Death” being the predictor and “Unlocking Phenotypic Plasticity” being the target). **G)** A scenario of having high relative importance and negative spatial dependency is illustrated in a bladder cancer sample (with “Deregulating Cellular Energetics” being the predictor and “Sustaining Proliferative Signaling” being the target).

Again, we explored the relationships between individual pairs of Hallmarks (**Figure 6D,E**). This revealed new patterns, such as the very high relative importance of “Resisting cell death” when predicting the spatial distribution of “Avoiding immune destruction” and “Tumor-promoting inflammation”, which has a negative spatial dependency. Another example is the particularly strong positive spatial dependency between “Evading growth suppressors” and “Activating invasion and metastasis”. This suggests cancer cells undergoing Evading growth suppressors are selectively predicting Activating invasion and metastasis process but not the other “cell-intrinsic” hallmarks particularly enabling replicative immortality and genome instability hallmarks. One possible interpretation of this result is that cancer cells that are fit to metastasize may be positively selected based on their ability to evade growth suppressors more so than having genomic instability or replicative immortality.

Similar to the previous experiment, we illustrated the relationship between a neoplastic Hallmark predicting its target TME Hallmark by overlaying the activities of a predictor-target pair on histology images from different cancer types. The case where a neoplastic Hallmark such as “Resisting Cell Death” has a high relative importance in predicting a target TME Hallmark such as “Unlocking Phenotypic Plasticity” through a positive spatial dependency is illustrated on a Kidney cancer sample (**Figure 6F**). By contrast, an example of a neoplastic Hallmark such as “Deregulating Cellular Energetics’’ having a high relative importance in predicting a target TME Hallmark such as “Sustaining Proliferative Signaling” through a negative spatial dependency is illustrated on a liver cancer sample (**Figure 6G**).

By faceting these results at the level of tumor types (**Figure S6B**), all but “(Nonmutational) Epigenetic Reprogramming” neoplastic Hallmarks seem to be equally important in predicting their targets through a mixed spatial dependency. In bladder cancer, however, “Resisting Cell Death” and “Deregulating Cellular Engergetics” best predict their targets likely through a mixed spatial dependency as well. Interestingly, “(Nonmutational) Epigenetic Reprogramming” seemed to have have higher importance. Another exceptional observation is in Glioblastoma, where the neoplastic Hallmark “Senescent cells” showed significant lower importance in predicting its targets compared to the rest of the cancer types.

## Discussion

If one were to summarize in two words the conceptual advances of the last decade in our understanding of oncogenesis, it might be “complexity” and “ecosystems”. On the one hand, the deep molecular characterization of thousands of human tumors through multiple molecular layers has revealed how dozens of key genes (oftentimes referred to as drivers) are driving different oncogenic processes. However, their overall functional consequences cannot be predicted through genomics alone, as their mutations have a complex interplay with each other, the tissue of origin of the tumor, other cells in the tumor microenvironment, the germline genome and clinical aspects of the patients, and a myriad of other factors. On the other hand, advances in single-cell genomics have shown how human tumors consist of an ever-increasing number of different cell types that, not only are co-existing within the tumor but, together, shape the phenotype and evolution of the tumor.

Cancer Hallmarks become a powerful tool in this context, as they simplify this molecular and cellular complexity to a set of principles or phenotypes that every tumor needs to achieve oncogenesis. This makes irrelevant whether such phenotypes are achieved through mutations in this or that pathway or assembled through the recruitment of this or that cell type (**Figure S5B**). The only thing that matters for the oncogenic process is whether the tumor achieves a specific activity of the Hallmark.

Here we exploited the concept of Cancer Hallmarks to simplify the complexity of our PanCancer spatial transcriptomics dataset and explore how these 13 critical phenotypes for oncogenesis are distributed through space across 63 samples of primary untreated tumors from 10 cancer types. We have presented a quantifiable approach demonstrating the spatial heterogeneity of Cancer Hallmarks across multiple cancer types and some of the key factors that drive such spatial distribution, namely cell identity, clonal divergence, and the spatial interdependence between the two major tumor compartments, cancer cells and the TME.

We first showed how each of the Cancer Hallmarks segregate into cancer cells or the TME. This near-binary classification of Hallmarks has been suspected, but here we demonstrate that it exists throughout 10 cancer types and follows a structured spatial pattern. This is important because, even if in Weinberg’s Cancer Hallmarks second iteration it was already stated that the TME plays an important role and can drive some of the Hallmarks^9^, it remained unclear which specific Hallmarks are contributed by each compartment, a question that we now answer, describe and quantify.

Furthermore, we also showed that the activity of Cancer Hallmarks within each compartment is also both variable and structured. The variability of gene expression in cancer cells has been recently explored at the single-cell level in a Pan-Cancer study^16^. Their results, suggesting that Cancer Hallmarks can be differentially active across cancer cells, agree with our observations of different areas within the neoplastic compartment having different Hallmark activities. However, we go one step further and show that such differences in Hallmark activities are spatially structured.

In the case of cancer cells, it is possible that differences in somatic mutations could partly drive this spatial structure: clones with different somatic mutations can also have different Cancer Hallmark activities. Indeed, our data suggest that cancer cells that diverge in genomic state tend to exhibit specialized Cancer Hallmark activities. However, we found this tendency to be relatively rare, as we only detected it in 3.39% of all pairs of clones. This agrees with recent results analyzing whole-genome sequencing with matching transcriptomics (at bulk level) from multiple samples from the same patients, showing that the expression of most genes in cancer cells is not inherited across clones^25^, but rather variable through other mechanisms. Therefore, even if genomic variability might lead to differential cancer Hallmark activity in several cases (i.e., specialization of some clones), this is often insufficient. Instead, our data suggest that cancer cells with different somatic mutations still need, in most cases, the activity of almost all Cancer Hallmarks. In the same study, the authors also suggest that interactions between cancer cells and the TME might be one of the main drivers of gene expression variability. Our results modeling the spatial interdependence between Hallmarks of cancer cells and those of the TME point towards that hypothesis. All Cancer Hallmarks from each compartment could be predicted with high accuracy using only the activity of the Hallmarks from the other compartment.

Based on these results, we propose a model of the overall architecture of Cancer Hallmarks in human primary tumors (**Figure 7**). In this model, cancer cells have different phenotypes depending on their relative position to the TME. The core of the neoplastic compartment (the area that is further from the TME) consists of cancer cells active in “Resisting Cell Death”, “Enabling Replicative Immortality”, “Genome Instability and Mutation” and “Deregulating Cellular Energetics”. On the other hand, at the interface with the TME we find cancer cells active in “Evading Growth Suppressors” and “Senescent cells” followed by “Non-Mutational Epigenetic Reprogramming” that’s closer to the core. In other words, we believe that there is a core of cancer cells (potentially cancer stem cells) that are exploring different phenotypic states through (epi)genetic alterations. This core is protected by a layer of cancer cells at the outer layers of the neoplastic compartment that is equipped with the phenotypes necessary to avoid any interference from the TME (**Figure 7A**).

**Figure 7.**
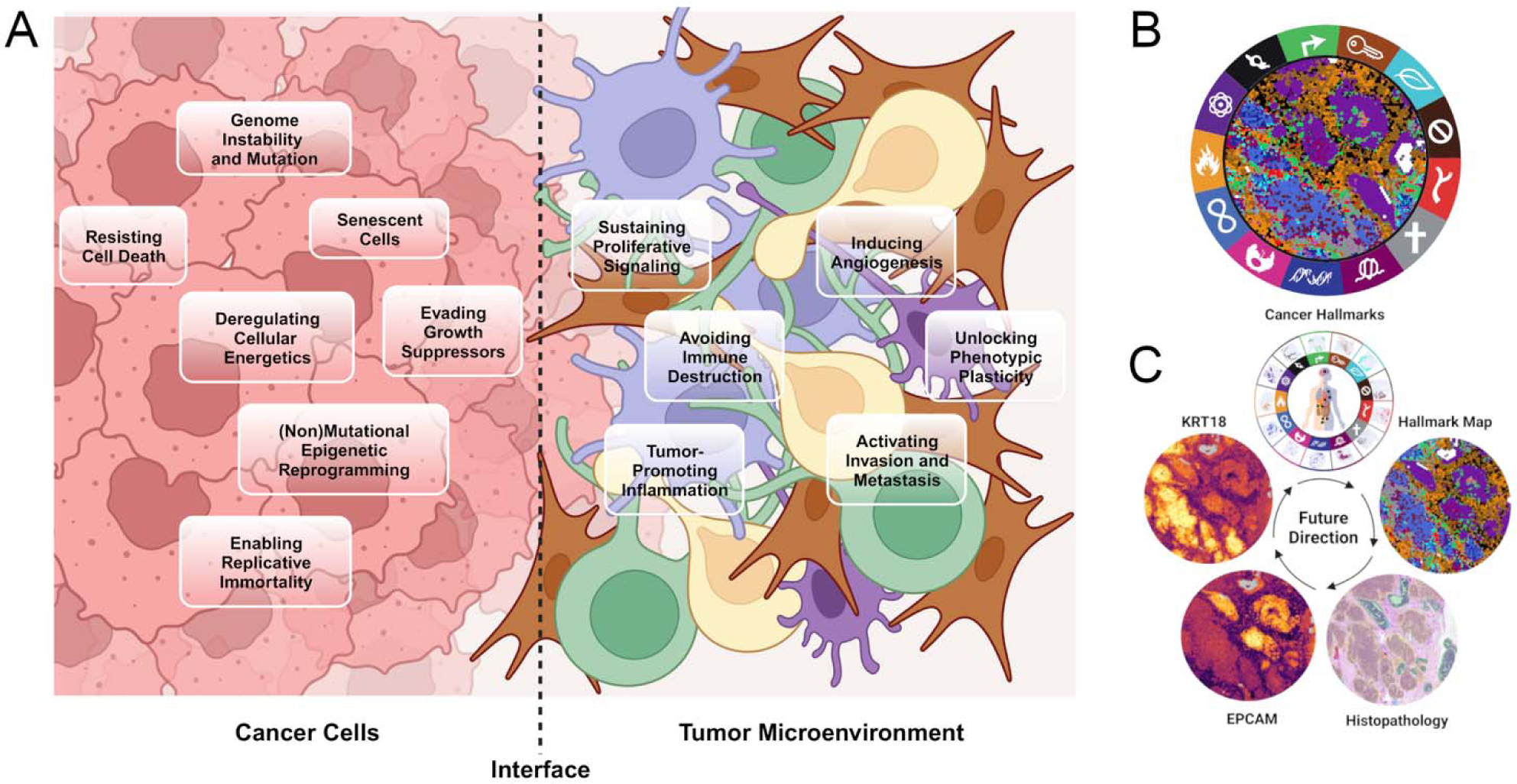
Implications of our work for basic and translational cancer research. **A)** Cancer cells may exhibit spatially structured divergent phenotypes for self-exploration (at the core) and in contact with the tumor microenvironment. The TME, on the other hand, collaborates through multiple phenotypes, at variable distances from the interface with cancer cells, to influence the spatial structure of the phenotypes of cancer cells. **B)** Hallmark Activity Map for one of the samples analyzed. Each VISIUM dot is colored according to its most active Hallmark. **C)** Schema depicting the potential applications and integration of our research within cancer pathology laboratories

From the TME perspective, while our data suggest most Hallmarks collaborate to configure the spatial landscape of Hallmarks of neoplastic cells (**Figure 5C**), they are spatially distributed depending on their relative position to the cancer cells. We find that at the interface with cancer cells there are cells that are active in “Sustaining Proliferative Signaling”, “Tumor-Promoting Inflammation”, and “Avoiding Immune Destruction”. We think the cells with the activity of these Hallmarks may play a dual role that generally would be to fight cancer cell proliferation, but then these same cells become subjugated by the cancer cells to promote self-sufficiency in growth signals and immune shut-down mechanisms. On the other hand, the core of the TME compartment (the area that is further from the collection of cancer cells) consists of cells highly active in “Activating Invasion and Metastasis”, “Inducing Angiogenesis”, and “Unlocking Phenotypic Plasticity”. We believe cancer cells leverage these core TME Hallmarks as a secondary mechanism after having conquered the immune system, eventually allowing them to invade nearby tissues and metastasize (**Figure 7A**).

This model is, of course, not necessarily universal, and it is likely to change across samples depending on other factors. For example, the existence of cancer clones that are specialized in specific Hallmarks due to genomic divergence might create a different architecture. Furthermore, we do not know at the moment whether cancer treatments alter the specifics of this model. In that sense, it is reasonable to hypothesize that immune-based therapies might lead to different outcomes depending on the initial spatial architecture of Hallmarks such as “Avoiding immune response” or “Resisting cell death”, among others.

From a translational perspective, we believe that the diversity in the spatial distribution of Cancer Hallmarks in the different solid tumors analyzed can help to understand the variability in the pharmacological and radiological responses found in the clinic when patients with molecular alterations considered drivers respond differently. Likewise, the prognostic role within each of the tumors that this spatial distribution may have is relevant, and creating spatial distribution signatures may contribute to a better understanding of the sensitivity to one or other treatment alternatives that may exist (**Figure 7B,C**). Definitely, these findings may contribute to improving how we will be able to stratify patients in future clinical trials.

In conclusion, this study shows how spatial transcriptomics data can be used to analyze the different Cancer Hallmarks, which describe how tumors behave and interact with their surroundings. We believe this approach has some advantages over methods such as purely single-cell-based analysis because it can show how the different features of the tumor are related to one another based on their physical location. However, future research should include a more detailed analysis of how our observed patterns that we observe relate to clinical outcomes. Overall, we believe that this approach has the potential to improve our understanding of cancer and ultimately lead to better treatment for patients.

## Limitations

Our spatial transcriptomics study comprised a unique dataset from 63 patients from 10 major tumor types. While being one of the largest spatial transcriptomics dataset to date, it is by no means exhaustive, in the sense that there are many cancer types that we have not been able to cover. Furthermore, we do not have any information about driver mutations or clinical outcomes of these patients, limiting our ability to extrapolate our findings to other molecular layers or clinical insights. Finally, another limitation stems from the fact that VISIUM does not provide single-cell resolution. Our study would have ideally benefited from a spatial transcriptomics technology at single-cell resolution for more accurate findings. Such technologies currently exist but are not yet as practical as VISIUM in several aspects, including the fact that they are limited to a few hundred genes, unlike the whole-transcriptome coverage offered by VISIUM. Nevertheless, we aimed to counteract this limitation by computationally enhancing the VISIUM resolution with the state-of-the-art BayesSpace method whenever possible.

## Acknowledgements

The authors would like to thank Abel Gonzalez-Perez, Nuria Lopez-Bigas, and Eliezer Van Allen for their valuable feedback. The authors would also like to acknowledge the kind and generous patients who donated the samples that made this study possible, as well as the scientists that generated the datasets for sharing them.

## Funding

M.S. and S.C. are funded by the Asociación Española Contra el Cáncer (AECC) project LABAE20038PORT. M.S. also received the support of a fellowship from “la Caixa” Foundation (LCF/BQ/DR22/11950021). E.P-P. is supported by a Ramon y Cajal fellowship from the Spanish Ministry of Science (RYC2019-026415-I), a Fundación FERO fellowship () and is a member of the 10x Clinical Translational Research Network. J. B. is supported by the Grupo Español de Investigación en Cáncer de Ovario (GEICO) Translational Research Award “Andrés Poveda” 2021. We thank CERCA Programme / Generalitat de Catalunya for institutional support. The research leading to these results has received funding from MCIN/ AEI /10.13039/501100011033/ and the European Development Regional Fund, ‘A way to make Europe’ ERDF (project PID2021-125282OB-I00); Departament de Recerca i Universitats / Generalitat de Catalunya (2021 SGR 01494); and the Sarah Jennifer Knott Foundation.

## Methods

### Data and code availability

The code used to run the analyses can be found in the following GitHub repository: https://github.com/CIG-spatialLandscape/spatialHallmarks_paper

### Generation of spatial transcriptomics data from fresh frozen samples

Frozen Ovarian Cancer samples (HGSOC) were embedded in cryomolds using OCT (Tissue Tek Sakura) using a bath of isopentane and liquid nitrogen (Tissue preparation guide CG000240, 10x Genomics). Following freezing in OCT, blocks were stored at −80°C. RNA integrity was assessed by calculating RNA Integrity Number (RIN) of freshly collected tissue sections with Qiagen protocol (RNeasy Mini Kit 74104) and analyzed by TapeStation. Samples with Rin above 7 were selected for the experiments. For sectioning, blocks were equilibrated to −20°C in the cryochamber (LEICA CM1950). 10 µm-thick sections were placed onto the active areas (6mm x 6mm) of chilled 10x genomics Visium slides. For Tissue Optimization, 7 sections were placed on a Tissue Optimization Slide (3000394, 10X Genomics) then fixed in chilled methanol and stained according to the Visium Spatial Tissue Optimization User Guide (<u>CG000238</u>, 10X Genomics) to determine optimal permeabilization time for HGSOC. For tissue optimization experiments, fluorescent images were taken with a TRITC filter using a 10X objective and 900 ms exposure time. For Gene Expression samples, 2 consecutive sections were placed on a chilled Visium Spatial Gene Expression Slide (2000233, 10X Genomics), and adhered by warming the back of the slide. Tissue sections were then fixed in chilled methanol and Hematoxilin & Eosin staining was performed according to the Demonstrated Protocol (CG000160, 10x genomics). Brightfield histology images were taken using a 10X objective (Plan APO) on a Nikon Eclipse Ti2, images were stitched together using NIS-Elements software (Nikon) and exported as tiff files. Following imaging, HGSOC samples were permeabilized for 30 minutes and cDNA Synthesis and amplification was performed following Visium Spatial Gene Expression User Guide (CG000239, 10X Genomics).

Libraries were prepared according to the Visium Spatial Gene Expression User Guide (CG000239, 10X Genomics) and sent for sequencing Using HiseqX 150PE (2x 150bp) applying 1% Phix. Sequencing depth was calculated with the formula (Coverage Area x total spots on the Capture Area) x 50,000 read pairs/spot. Sequencing was performed using the following read protocol: read 1: 28 cycles; i7 index read: 10 cycles; i5 index read: 10 cycles; read 2: 90 cycles.

### Generation of spatial transcriptomics data from formalin-fixed paraffin-embedded (FFPE) samples

RNA integrity was assessed by calculating DV200 of RNA extracted from freshly collected tissue sections. Briefly, Tissue blocks were placed in the microtome and cut to expose the tissue. 4 sections 10 µm thick were placed in a chilled Eppendorf tube and RNA extraction protocol from Qiagen was performed (Rneasy FFPE Kit 73504) and RNA was analyzed by TapeStation. 21 Samples with DV200 ≥ 25% were selected for experiments. FFPE samples were placed in the microtome and sectioned 7 µm thick, after floating on a water bath at 42 °C, sections were placed on Visium Spatial Gene Expression slides (2000233, 10X Genomics). After sectioning, the slides were dried at 42°C for 3 hours. The slides were then placed inside a slide mailer, sealed with parafilm, and left overnight at Room temperature. The slides were deparaffinized by successive immersions in xylene and ethanol followed by H&E staining according to Demonstrated Protocol (CG000409, 10X Genomics).

Brightfield images were taken using a 10X objective (Plan APO) on a Nikon Eclipse Ti2, images were stitched together using NIS-Elements software (Nikon) and exported as tiff files. After imaging, the glycerol and cover glass were carefully removed from the Visium slides by holding the slides in an 800 ml water beaker and letting the glycerol diffuse until the cover glass detached and density changes were no longer visible in the water. The slides were then dried at 37°C.

Libraries were prepared according to the Visium Spatial Gene Expression for FFPE User Guide (CG000407, 10X Genomics) and sent for sequencing Using HiseqX 150PE (2x 150bp) applying 1% Phix. Sequencing was performed using the specific for FFPE following read protocol: read 1: 28 cycles; i7 index read: 10 cycles; i5 index read: 10 cycles; read 2: 50 cycles.

### Sequencing, data processing and filtering

Raw fastq files of two ovarian cancer, four colorectal cancer, four breast cancer, five bladder cancer, five prostate cancer, seven lung cancer, and 4 glioblastoma samples were processed with the “spaceranger” command line tool (version 1.3.1, 10x Genomics) and mapped to the pre-built human reference genome (GRCh38). Thirty-six publicly available VISIUM tumor datasets from seven tissues were downloaded from recent publications^19, 28–34^ and the ‘spatial gene expression’ panel of 10x Genomics (https://www.10xgenomics.com/resources/datasets). The samples were obtained from the following sections: ‘Visium Spatial Targeted Demonstration Data (v1)’, ‘Visium Spatial Fluorescent Demonstration (v1)’, ‘Visium Spatial for FFPE Demonstration (v1)’. For each downloaded ST dataset, the output from SpaceRanger was loaded individually in the STutility environment^35^. Each sample was filtered following three steps. First, genes with less than 5 UMIs were excluded. Second, mitochondrial and ribosomal genes, and non protein-coding genes were removed from the analysis. Genes that were kept were tagged as protein_coding, TR_V_gene, TR_D_gene, TR_J_gene, TR_C_gene, IG_LV_gene, IG_V_gene, IG_J_gene, IG_C_gene and IG_D_gene. Lastly, spots were filtered out depending on two conditions: if they were isolated from the majority of the tissue and/or if the number of features was lower than a threshold value established by the distribution in each sample. Each filtered sample was then individually normalized by variance stabilizing transformation on the expression data using SCTransform (return.only.var.genes = FALSE, variable.features.n = NULL, variable.features.rv.th = 1.1)^36^.

### Spatial enhancement and gene expression imputation with BayesSpace

As each of the VISIUM spots may contain a number of cells (1-10), we sought to conduct our downstream analysis in a way that better approximates single-cell resolution. We therefore used the novel BayesSpace R package that leverages a fully Bayesian statistical framework by using information from spatial neighborhoods to computationally enhance the resolution of spatial transcriptomics data^21^. To that end, the normalized gene expression data was converted to a SingleCellExperiment object with the respective spatial coordinates and metadata. In the preprocessing step (spatialPreprocess), normalization was skipped and 15 principal components (PCs) were computed using the top 2000 highly variable genes in the SCT assay. The number of clusters was chosen by the elbow of the pseudo-log-likelihood plot (qTune). Then, spots were clustered by spatialCluster using 50000 iterations, 15 PCs, gamma = 3 (recommended for Visium), and the initialization method k-means. To increase the resolution of the data, each spot was divided into 6 subspots using spatialEnhance from BayesSpace. For those samples that were processed using the “-reorient-images” flag in SpaceRanger, their “imagerow” and “imagecol” columns were switched to prevent inaccurate enhanced clustering. The resulting clusters were used as the initialization of the algorithm using 200000 iterations and 0.3 of prior jitter. Finally, xgboost was used to impute the expression of the complete set of genes for each subspot with a maximum number of boosting iterations of 100.

### Assigning genes to Hallmarks

The full list of pathways was downloaded from the Pathway Commons data portal (v12, 2019) (http://www.pathwaycommons.org/archives/PC2/v12/). From this collection, pathways with less than 4 genes or more than 475 genes in their corresponding gene sets were discarded, as were those with “CHEBI” symbol corresponding to small chemical compounds, yielding a total of 3,250 pathways. To assign genes for each Hallmark, a two-stage pathway filtering process was undertaken. First, specific keywords that can be associated with specific Hallmarks were searched in the names of the pathways following a similar previously reported approach^46^, and the resulting pathways were assigned to their associated Hallmark. Second, for Hallmarks that were assigned an excessive number of pathways, ChatGPT (v. 3.5) was used to narrow-down the number of pathways in those Hallmarks by prompting it to retain pathways that are minimally involved across Hallmarks. Following this pathway-to-Hallmark assignment, each list was manually inspected to ensure the credibility of these assignments. This approach was applied for all 13 Hallmarks except “Resisting Cell Death” as it required a more careful assignment of pathways. Since pathways that are directly related to cell death mechanisms such as apoptosis may contain pro-as well as anti-apoptotic genes, this could mask the downstream signature interpretation. We therefore manually assigned a number of pathways that could directly be related to survival mechanisms, which would indirectly imply resisting cell death.

### Assigning the neoplastic and TME compartments

“ESTIMATE” R package was used to quantify the neoplastic cell purity within each subspot in the enhanced VISIUM dataset^22^. This package has been extensively and successfully used to quantify the presence of cancer, immune or stromal cells in RNAseq profiles from bulk tumor samples. As such, we assumed that, since each (sub)spot in our dataset can contain multiple cells (often between 1 and 10), it could be treated as a mini-bulk RNAseq and, therefore, deconvoluted with the same approach. To perform the quantification, the gene expression of each subspot was first extracted and transformed into GCT format. Then, the “estimate” function was used to compute the stroma, immune, and ESTIMATE scores. Finally, ESTIMATE scores were added as metadata, and they were used to create five clusters within each sample using the k-means algorithm. The cluster with the lowest ESTIMATE values corresponds to the maximum purity of neoplastic cells, whereas the cluster with the highest ESTIMATE values corresponded to the minimum purity of neoplastic cells (TME). An expert pathologist validated these scores, who determined the neoplastic and stroma cell content in each cluster through H&E images from seventeen samples.

### Scoring the Hallmarks

To compute the activity of a Hallmark, the enhanced object was first converted back to Seurat, and the expression matrix was scaled (ScaleData) to perform downstream analysis. A module score was computed for each Hallmark’s gene set using AddModuleScore from Seurat^36^ on the enhanced objects, followed by scaling and centering the obtained scores within each sample, yielding what we refer to as “Hallmark activity”. The resulting Hallmark activities were then averaged per ESTIMATE cluster depending on the neoplastic cell content. Rows and columns were clustered using “average” as the clustering method and “correlation” as the distance measure after scaling the columns. The same Seurat function was used to score the activities of each individual pathway from which the genes formed the signature of “(Non)mutational Epigenetic Reprogramming”.

### Building Gene Exclusion Controls

Two levels of controls were built in order to ensure the reliability of the genes in the selected pathways to sufficiently represent the quantified Hallmark scores. First, 25% of the genes from each Hallmark-associated pathway were randomly excluded and the remaining genes were combined for each Hallmark to form a control gene signature. The second level was similarly applied with 50% exclusion. Each level was repeated 5 times to obtain a total of 10 randomized Hallmark control signatures. The controls were scored using the same parameters with AddModuleScore from Seurat.

### Building Gene Addition Controls

In order to investigate whether adding new genes to each of the 13 Hallmark gene sets could influence the signature scores, the percentage of overlap between the genes of each pathway (excluding the final 109 pathways) from the initially filtered list (3,250 pathways) and the genes of each of the 13 Hallmark gene-sets was computed. Pathways with an overlap of at least 60% and at most 90% were subsequently assigned as an accessory pathway list for the associated Hallmark based on an assumption that those pathways with such a high overlap and which were not included initially may also be important for the associated Hallmark. The union of the unique set of genes from these accessory pathway lists associated with one Hallmark were stored as a list of accessory genes that could be assigned to that Hallmark. From this accessory list for each Hallmark, a number of genes corresponding to 5%, 10%, 25%, and 50% of the size of the original Hallmark-associated gene set were selected and added to the original gene set. This process was repeated 5 times for each of the added percentages. The resulting Hallmark gene set controls were scored using the same parameters with AddModuleScore from Seurat.

### Correction of the distances between subspots

The distance between the center of two adjacent spots is 100 μm in the VISIUM technology, which has a spot diameter of 55 μm. These distances must be considered after enhancing, as two subspots coming from the same spot will have a shorter distance than two adjacent subspots from two different spots. Therefore, the spot location grid created by BayesSpace at the spot level was multiplied by 100 μm to preserve the distance between spots. Then, each subspot was shifted from the center following the same hexagonal distribution as BayesSpace enhancing, but instead of applying 1/3 of the distance between spots, a third of the radius (27.5/3) was used to determine the center of each subspot.

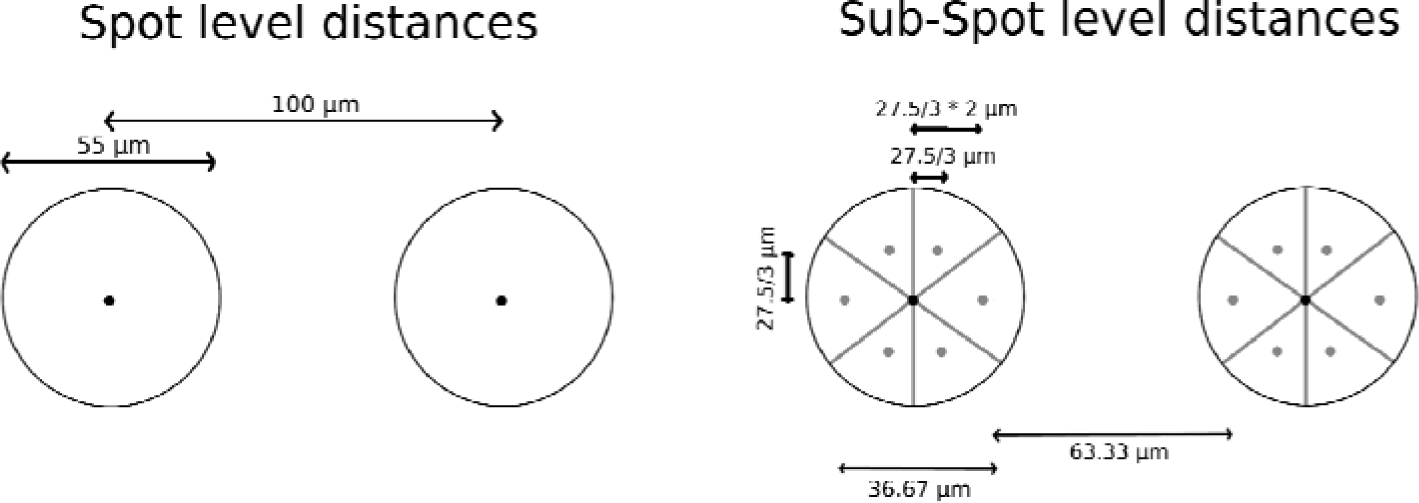

### Spatial autocorrelation of Hallmark activities

To determine the spatial autocorrelation of Hallmark activities in the tissue, we computed Moran’s I metric by modifying the function CorSpatialGenes() from STutility to apply it to our enhanced VISIUM samples. Each neighborhood was computed using 10 neighbors (5 subspots coming from the same spot and 5 others from adjacent spots) and weighted by the corrected subspot distances. Moran’s I was computed for the 7 TME Hallmarks within ESTIMATE clusters 4&5 and the 6 neoplastic Hallmarks within ESTIMATE clusters 1&2, and for all 13 Hallmarks within the whole tissue of each sample.

### Building Spatial Distribution Controls

Control signatures were built to compare their spatial autocorrelation with those of Hallmark signatures. These controls were generated by randomly selecting genes of various sizes: 50, 100, 200, 350, and 500. For each size, 25 sets of genes were randomly sampled from the intersection of all genes captured across all samples. The controls were scored using the same parameters with AddModuleScore from Seurat. Subsequently, Moran’s I was calculated for each gene set, and the average value was determined for each sample and size.

### Detection of cancer clones by quantifying copy number alterations

inferCNV method was used to detect copy number alterations (CNA) in each spot based on gene expre sion. Initially, the compartment labels from ESTIMATE clusters were assigned to the spot resolution using the max-voting rule. In case of a tie, the label “Buffer” was assigned. Subsequently, raw gene expression counts were used to infer CNA in neoplastic spots (clusters 1 & 2) with TME spots (clusters 4 & 5) serving as a reference. The “Buffer” spots were excluded from this analysis. The inferCNV function was executed with the following parameters: run(object_infCNV, num_threads = 1, cutoff=0.1, cluster_by_groups=F, plot_steps=T, denoise=T, no_prelim_plot=F, k_obs_groups = 1, HMM=T, leiden_resolution = 0.005).

The output results were obtained from step 17, which included spot groups from the dendrogram and the CNA state of each gene. We then transferred back to the subspot resolution, excluding spot groups with fewer than 10 spots (60 subspots) as they might be outliers in the dendrogram. Next, we calculated the genomic distance between each pair of groups within a sample by determining the number of genes with the same CNA state divided by the total number of genes used to infer CNA states (JSI score). To quantify the Hallmark difference, we averaged the Hallmark score within each group and computed the absolute difference. Finally, we correlated, for each sample and Hallmark, the genomic distance (1 -JSI) and the absolute Hallmark difference using Pearson correlation.

### Modeling the spatial relationship between Cancer Hallmarks of the neoplastic and those of the TME compartments with random forests

Before running the machine learning experiments, the scaled Hallmark scores were shifted to positive values in order to minimize the effect of negative values from the originally scaled Hallmark scores on downstream interpretation. The shifted scores were calculated by transforming the minimum scaled Hallmark scores into zero and shifting the rest of the values accordingly. A small constant is then added for each value to prevent dividing by zero in downstream analysis. These shifted values are then used in the input of the following machine learning experiments.

The spatial relationship between the Hallmarks of the neoplastic (ESTIMATE clusters 1&2) and TME (ESTIMATE clusters 4&5) compartments was modeled by running two machine learning experiments:

1. predicting the spatial distribution of each one of the 7 neoplastic Hallmarks as individual targets using the 6 TME Hallmarks altogether as the features (predictors), yielding 7 targets * 63 samples = 441 models in total.
2. vice-versa: predicting the spatial distribution of each one of the 6 TME Hallmarks as individual targets using the 7 neoplastic Hallmarks altogether as the features (predictors), yielding 6 targets * 63 samples = 378 models in total.

In either of the experiments, as the predictors must be located in the same subspots (compartment) where the target is located, we therefore translated the activity of predictors from their original compartment into an activity within the target compartment that serves much like a “radar” (**Figure 4A & S4**). These “radar” scores are constructed in each of the 63 samples that ultimately serve as the predictors in each model. In the case of the first experiment, the radar scores for each of the 6 TME Hallmarks in each neoplastic subspot are calculated following 2 major steps:

1. Each “radar” in each neoplastic subspot first captures the shifted Hallmark score from each one of the TME subspots weighted by the inverse of the corrected distance between each TME subspot and the neoplastic subspot where the radar is located.
2. These individual spatially weighted scores that are coming from all TME subspots in each neoplastic subspot are then summed up into a single score for each neoplastic subspot, which we finally call the “radar” score.

The same process is applied reversely when running the second experiment. Radar scores as predictors of their target Hallmarks from either experiments are then used to train the model on 80% of randomly-selected subspots with the ranger package^37^, followed by predicting the target Hallmark activity (response) using the rest 20% of the subspots with the computation of the shapley additive values (SHAP)^38^. We used the SHAP values to identify how the model prioritizes the relationships between the features (radars) and the target Hallmark using two metrics. The first is feature importance, which is the mean of absolute values of SHAP values. After scaling these values across features from 0 to 100, the fraction of each feature is used to infer the relative importance of that feature in predicting its target Hallmark. The second metric is feature dependency, which profiles the relationship between the feature and its SHAP value as a proxy to its target. Pearson correlation is calculated between them to infer the directionality of these relationships. Finally, as the predictors are weighted by distances we, therefore, call the latter metric in this context as “spatial dependency” for a more intuitive understanding.

### TME cell type composition

Cell type markers from the *PangloaDB* database^39^ were used to quantify the spatial distribution of certain cell types pertinent to the TME. Cell types from three main organ categories were selected from the database to represent the TME; the immune system, vasculature, and the connective tissue (only Fibroblasts). For each cell type, only canonical markers were obtained, followed by retaining cell types with at least 10 genes, yielding a total of 23 cell types (**Table S6**).

### Graphical figures

Individual panels with statistical data were generated with the R packages “ggplot2”, “pheatmap”, “ComplexHeatmap”. Figures 3A, 4A, S4A, and 7 were generated using BioRender, and all individual panels were put together using GIMP.

## Notes

### Competing Interest Statement

E.P-P is part of the Clinical Translational Research Network sponsored by 10x Genomics

### Summary of Updates

We have updated several analyses and results following various reviewer's comments

