## Supplementary Figures for "The spatial landscape of Cancer Hallmarks reveals patterns of tumor ecology"

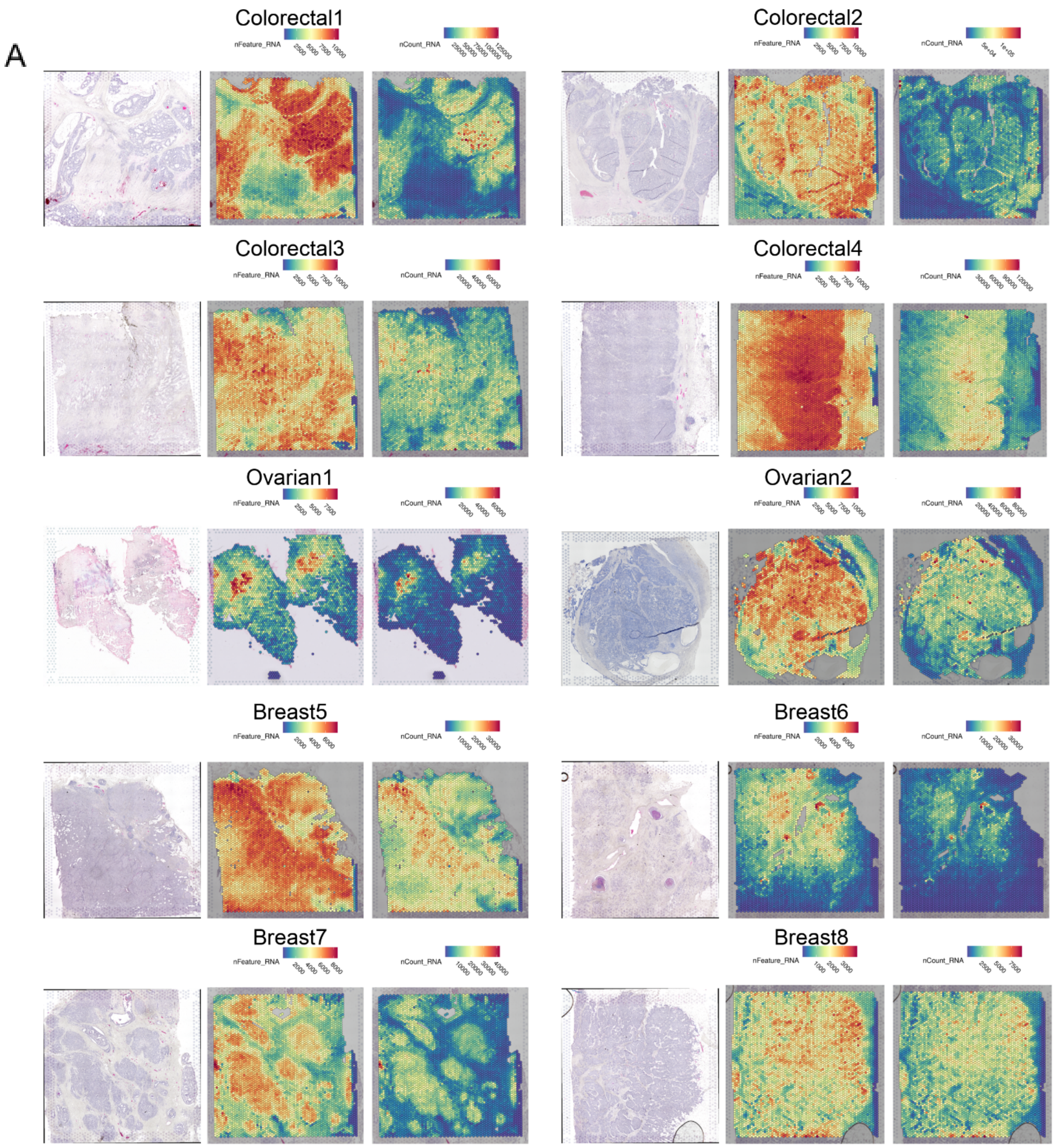
A
Colorectal1
Colorectal2
Colorectal3
Colorectal4
Ovarian1
Ovarian2
Breast5
Breast6
Breast7
Breast8
nFeature_RNA
nCount_RNA

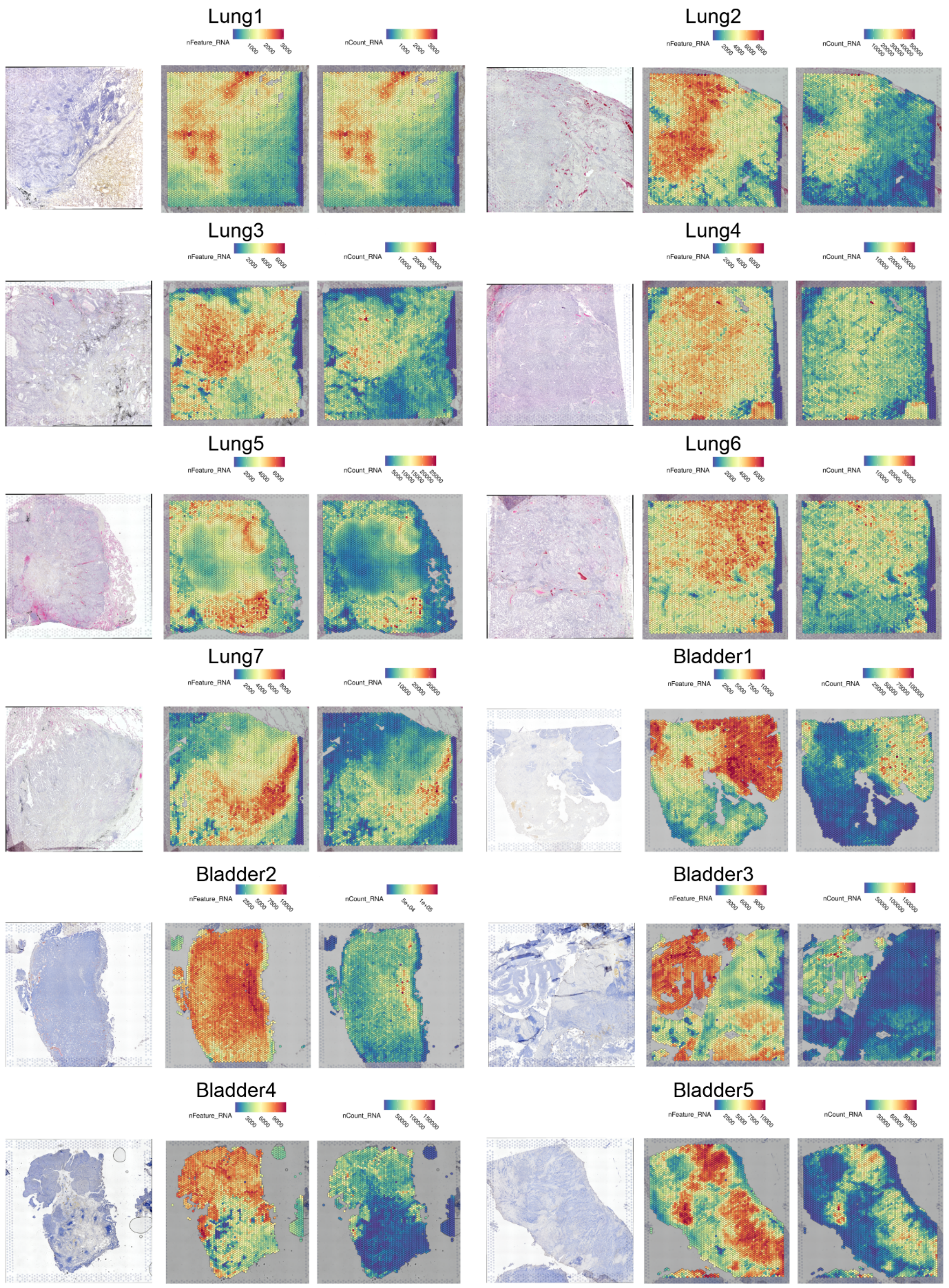
Lung1
nFeature_RNA 1000 2000 3000
nCount_RNA 1000 2000 3000
Lung2
nFeature_RNA 2000 4000 6000 8000
nCount_RNA 10000 20000 30000 40000 50000
Lung3
nFeature_RNA 2000 4000 6000
nCount_RNA 10000 20000 30000
Lung4
nFeature_RNA 2000 4000 6000
nCount_RNA 10000 20000 30000
Lung5
nFeature_RNA 2000 4000 6000
nCount_RNA 5000 10000 15000 20000 25000
Lung6
nFeature_RNA 2000 4000 6000
nCount_RNA 10000 20000 30000
Lung7
nFeature_RNA 2000 4000 6000 8000
nCount_RNA 10000 20000 30000
Bladder1
nFeature_RNA 2500 5000 7500 10000
nCount_RNA 25000 50000 75000 100000
Bladder2
nFeature_RNA 2500 5000 7500 10000
nCount_RNA 5e+04 1e+05
Bladder3
nFeature_RNA 3000 6000 9000
nCount_RNA 50000 100000 150000
Bladder4
nFeature_RNA 3000 6000 9000
nCount_RNA 50000 100000 150000
Bladder5
nFeature_RNA 2500 5000 7500 10000
nCount_RNA 30000 60000 90000

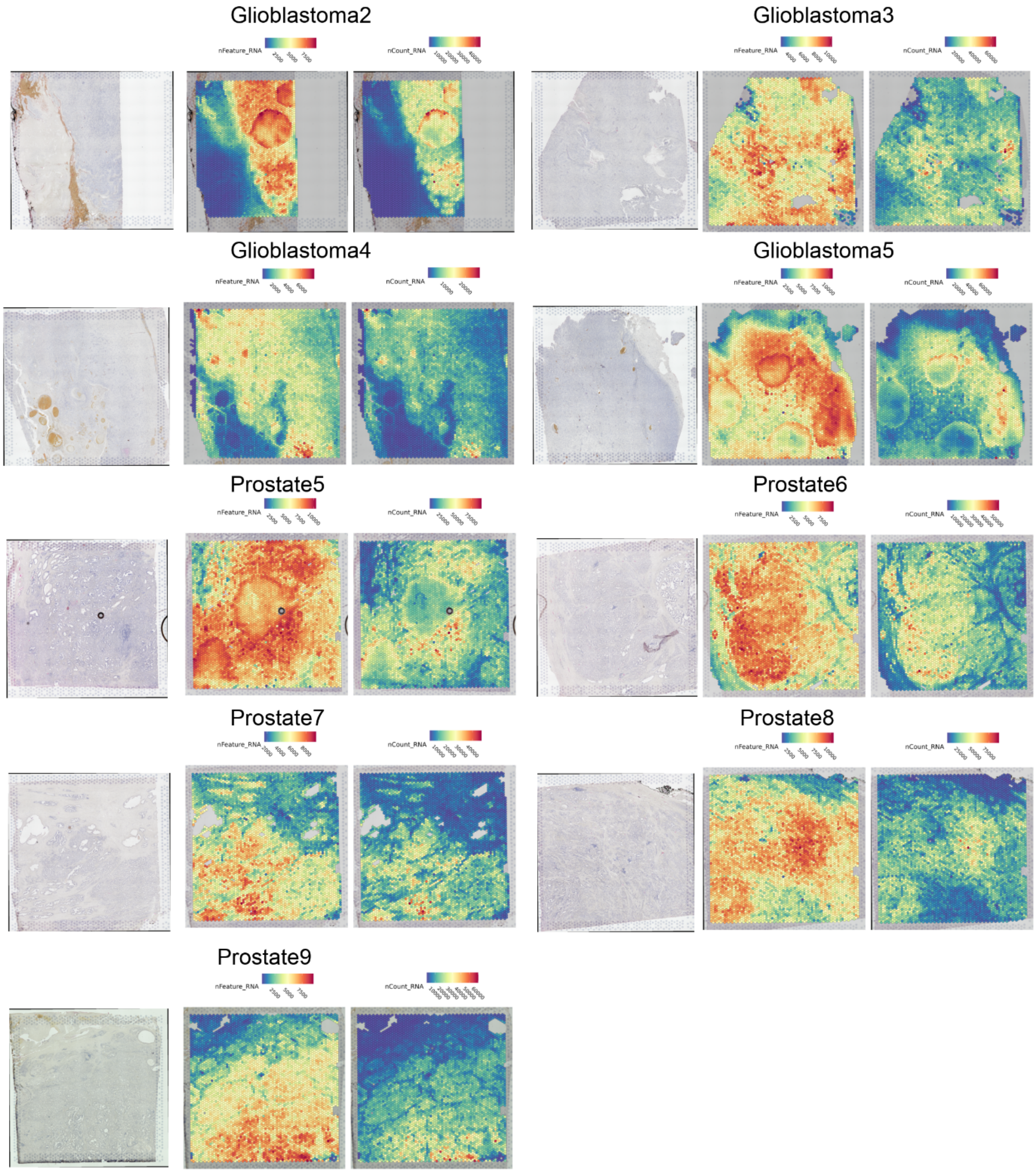
Glioblastoma2
nFeature_RNA
nCount_RNA
Glioblastoma3
nFeature_RNA
nCount_RNA
Glioblastoma4
nFeature_RNA
nCount_RNA
Glioblastoma5
nFeature_RNA
nCount_RNA
Prostate5
nFeature_RNA
nCount_RNA
Prostate6
nFeature_RNA
nCount_RNA
Prostate7
nFeature_RNA
nCount_RNA
Prostate8
nFeature_RNA
nCount_RNA
Prostate9
nFeature_RNA
nCount_RNA

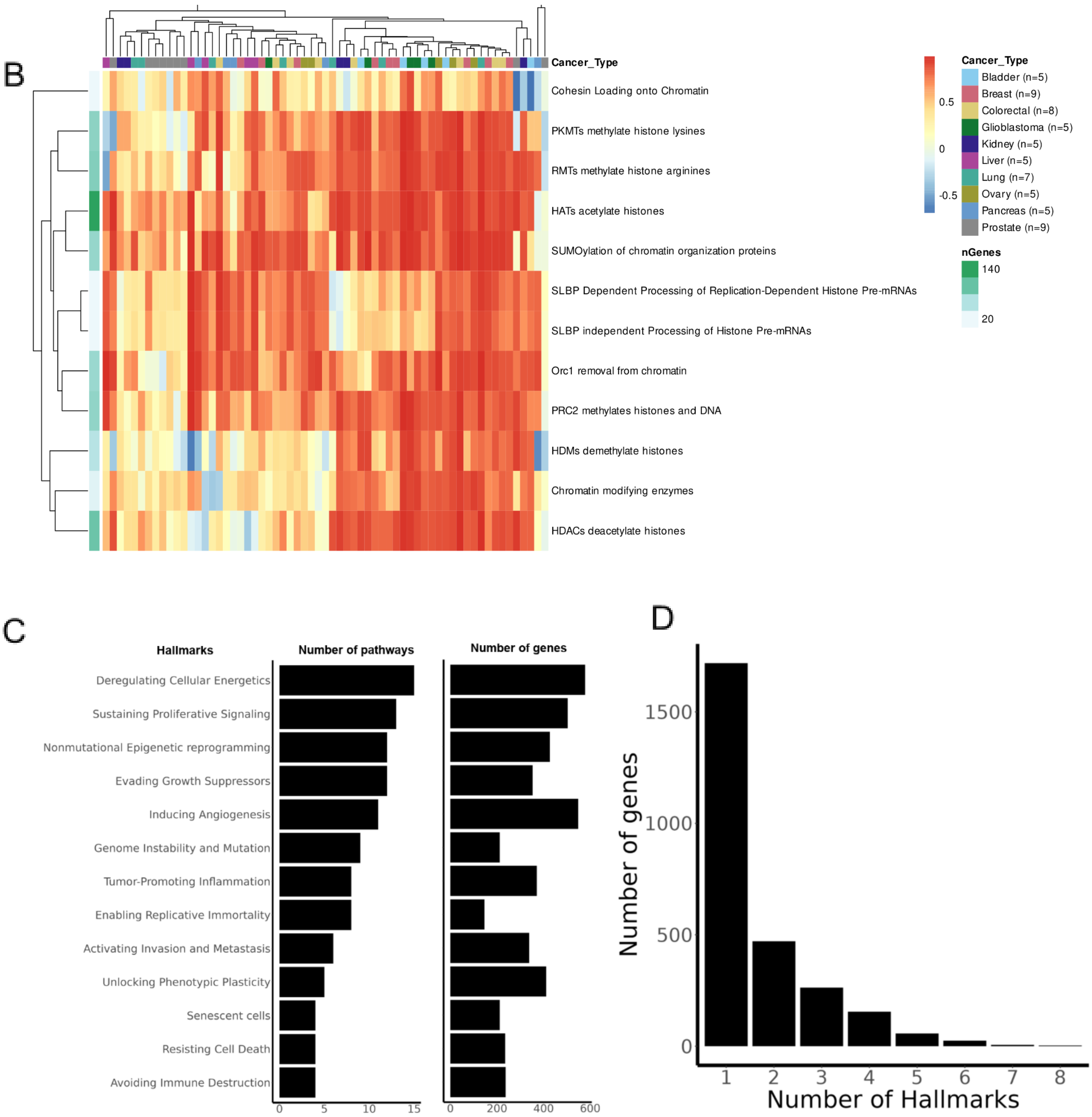

B
Cancer_Type
Cohesin Loading onto Chromatin
PKMTs methylate histone lysines
RMTs methylate histone arginines
HATs acetylate histones
SUMOylation of chromatin organization proteins
SLBP Dependent Processing of Replication-Dependent Histone Pre-mRNAs
SLBP independent Processing of Histone Pre-mRNAs
Orc1 removal from chromatin
PRC2 methylates histones and DNA
HDMs demethylate histones
Chromatin modifying enzymes
HDACs deacetylate histones
0.5
0
-0.5
Cancer_Type
Bladder (n=5)
Breast (n=9)
Colorectal (n=8)
Glioblastoma (n=5)
Kidney (n=5)
Liver (n=5)
Lung (n=7)
Ovary (n=5)
Pancreas (n=5)
Prostate (n=9)
nGenes
140
20
C
Hallmarks
Number of pathways
Number of genes
Deregulating Cellular Energetics
Sustaining Proliferative Signaling
Nonmutational Epigenetic reprogramming
Evading Growth Suppressors
Inducing Angiogenesis
Genome Instability and Mutation
Tumor-Promoting Inflammation
Enabling Replicative Immortality
Activating Invasion and Metastasis
Unlocking Phenotypic Plasticity
Senescent cells
Resisting Cell Death
Avoiding Immune Destruction
0
5
10
15
0
200
400
600
D
Number of genes
1500
1000
500
0
1
2
3
4
5
6
7
8
Number of Hallmarks

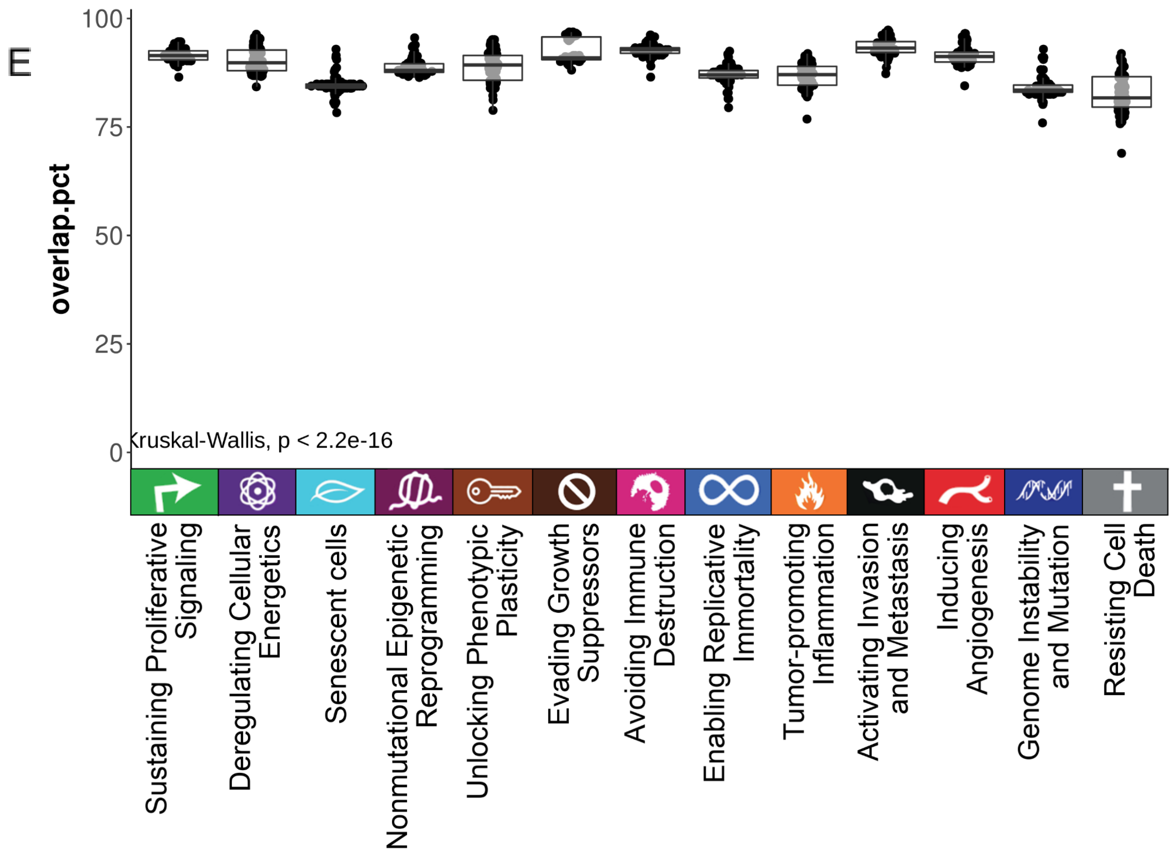
E
100
75
50
25
0
overlap.pct
Kruskal-Wallis, p < 2.2e-16
Sustaining Proliferative Signaling
Deregulating Cellular Energetics
Senescent cells
Nonmutational Epigenetic Reprogramming
Unlocking Phenotypic Plasticity
Evading Growth Suppressors
Avoiding Immune Destruction
Enabling Replicative Immortality
Tumor-promoting Inflammation
Activating Invasion and Metastasis
Inducing Angiogenesis
Genome Instability and Mutation
Resisting Cell Death

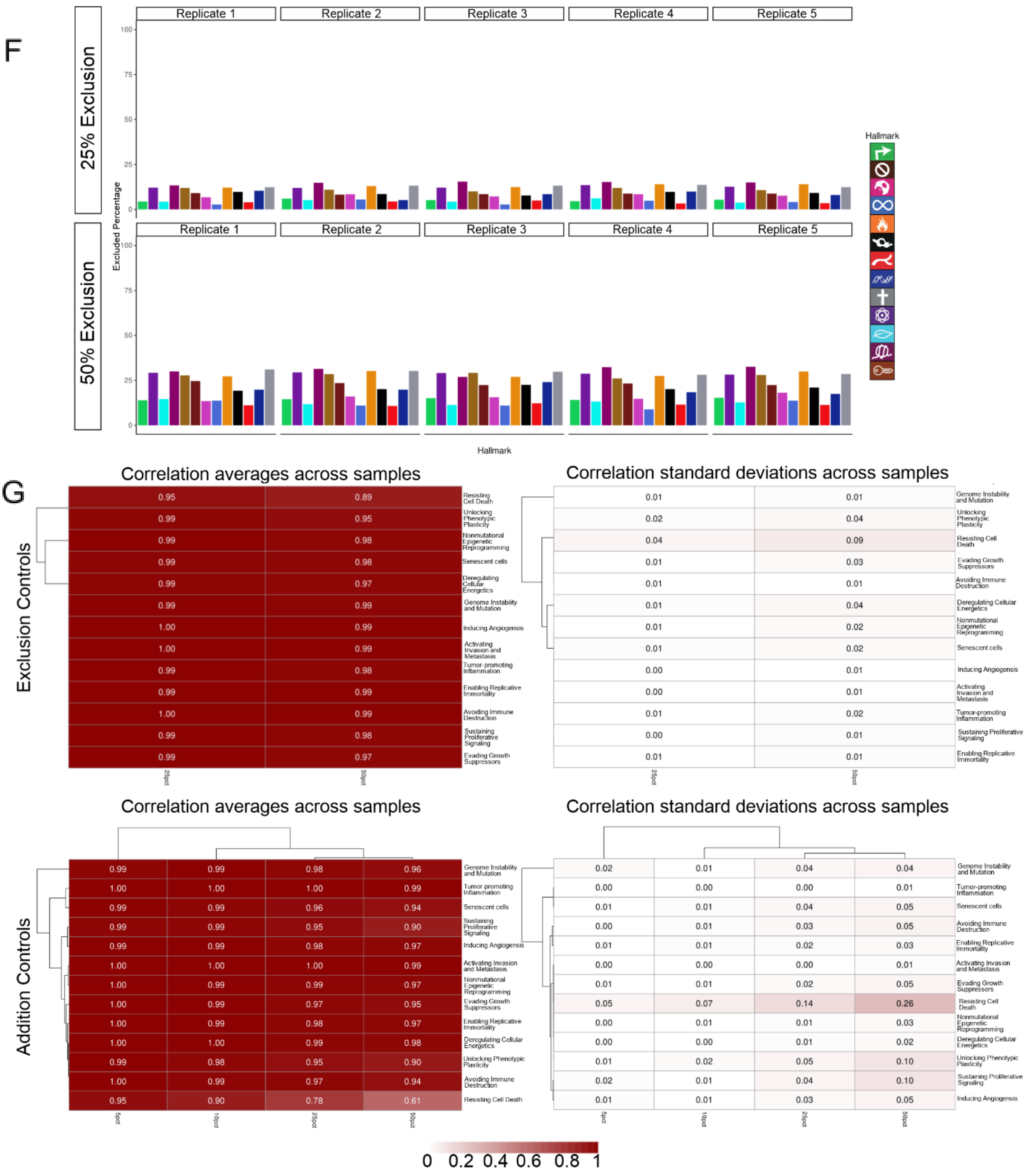
F
25% Exclusion
50% Exclusion
Replicate 1
Replicate 2
Replicate 3
Replicate 4
Replicate 5
Excluded Percentage
100
75
50
25
0
Hallmark
Hallmark
G
Correlation averages across samples
Exclusion Controls
Resisting Cell Death 0.95 0.89
Unlocking Phenotypic Plasticity 0.99 0.95
Nonmutational Epigenetic Reprogramming 0.99 0.98
Senescent cells 0.99 0.98
Deregulating Cellular Energetics 0.99 0.97
Genome Instability and Mutation 0.99 0.99
Inducing Angiogensis 1.00 0.99
Activating Invasion and Metastasis 1.00 0.99
Tumor promoting Inflammation 0.99 0.98
Enabling Replicative Immortality 0.99 0.99
Avoiding Immune Destruction 1.00 0.99
Sustaining Proliferative Signaling 0.99 0.98
Evading Growth Suppressors 0.99 0.97
25pct
50pct
Correlation standard deviations across samples
Genome Instability and Mutation 0.01 0.01
Unlocking Phenotypic Plasticity 0.02 0.04
Resisting Cell Death 0.04 0.09
Evading Growth Suppressors 0.01 0.03
Avoiding Immune Destruction 0.01 0.01
Deregulating Cellular Energetics 0.01 0.04
Nonmutational Epigenetic Reprogramming 0.01 0.02
Senescent cells 0.01 0.02
Inducing Angiogensis 0.00 0.01
Activating Invasion and Metastasis 0.00 0.01
Tumor-promoting Inflammation 0.01 0.02
Sustaining Proliferative Signaling 0.00 0.01
Enabling Replicative Immortality 0.01 0.01
25pct
50pct
Correlation averages across samples
Addition Controls
Genome Instability and Mutation 0.99 0.99 0.98 0.96
Tumor promoting Inflammation 1.00 1.00 1.00 0.99
Senescent cells 0.99 0.99 0.96 0.94
Sustaining Proliferative Signaling 0.99 0.99 0.95 0.90
Inducing Angiogensis 0.99 0.99 0.98 0.97
Activating Invasion and Metastasis 1.00 1.00 1.00 0.99
Nonmutational Epigenetic Reprogramming 1.00 0.99 0.99 0.97
Evading Growth Suppressors 1.00 0.99 0.97 0.95
Enabling Replicative Immortality 1.00 0.99 0.98 0.97
Deregulating Cellular Energetics 1.00 1.00 0.99 0.98
Unlocking Phenotypic Plasticity 0.99 0.98 0.95 0.90
Avoiding Immune Destruction 1.00 0.99 0.97 0.94
Resisting Cell Death 0.95 0.90 0.78 0.61
5pct
10pct
25pct
50pct
Correlation standard deviations across samples
Genome Instability and Mutation 0.02 0.01 0.04 0.04
Tumor-promoting Inflammation 0.00 0.00 0.00 0.01
Senescent cells 0.01 0.01 0.04 0.05
Avoiding Immune Destruction 0.00 0.01 0.03 0.05
Enabling Replicative Immortality 0.01 0.01 0.02 0.03
Activating Invasion and Metastasis 0.00 0.00 0.00 0.01
Evading Growth Suppressors 0.01 0.01 0.02 0.05
Resisting Cell Death 0.05 0.07 0.14 0.26
Nonmutational Epigenetic Reprogramming 0.00 0.01 0.01 0.03
Deregulating Cellular Energetics 0.00 0.00 0.01 0.02
Unlocking Phenotypic Plasticity 0.01 0.02 0.05 0.10
Sustaining Proliferative Signaling 0.02 0.01 0.04 0.10
Inducing Angiogensis 0.01 0.01 0.03 0.05
5pct
10pct
25pct
50pct
0 0.2 0.4 0.6 0.8 1

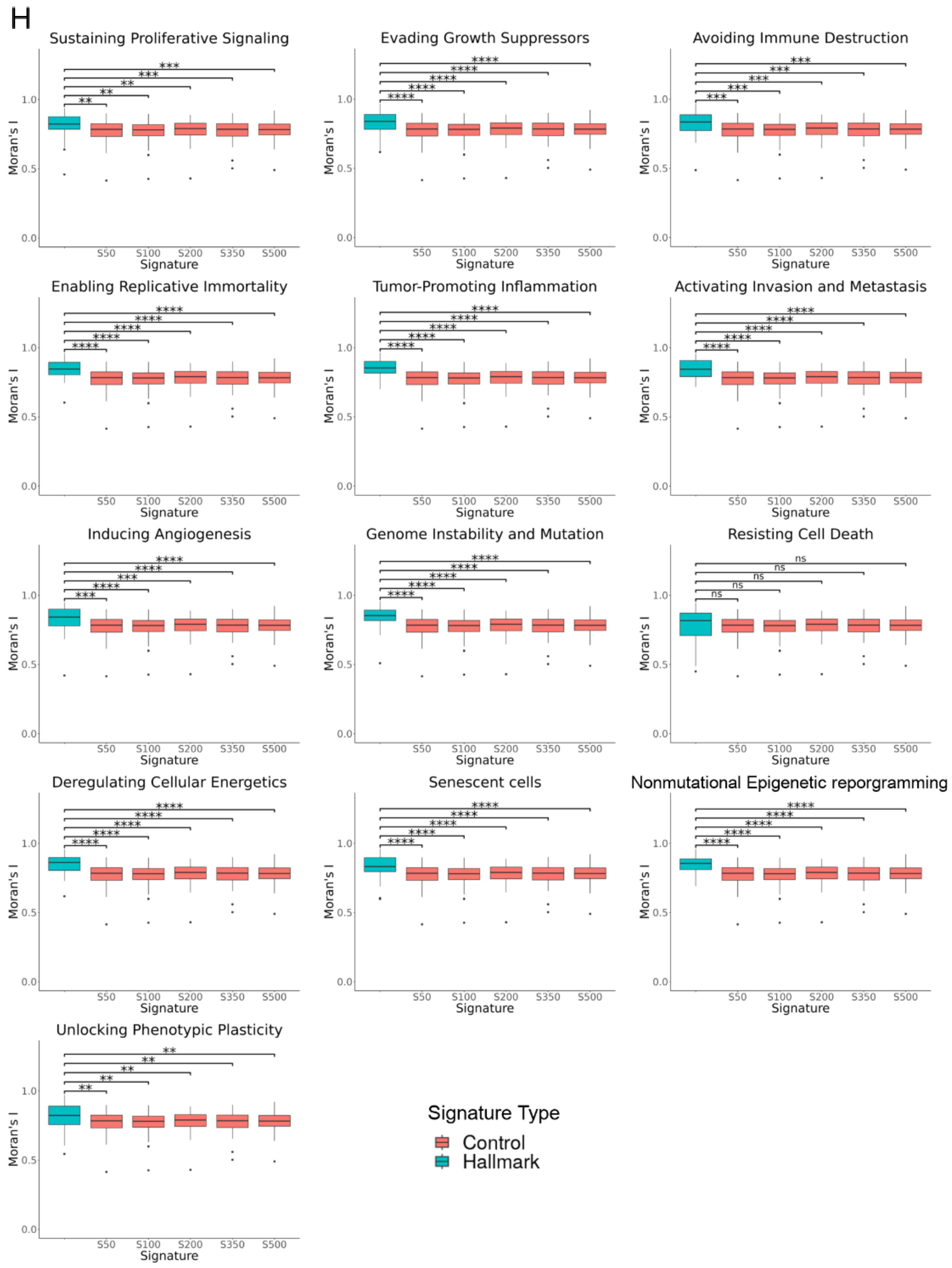


**Figure S1: Descriptions of quality metrics, assignments of Hallmark-associated genes and various controls for Hallmark signatures. A)** H&E images, number of unique genes and the number of UMIs per spot for each of the 31 samples that was generated in-house. **B)** Heatmap depicts the average Pearson correlation coefficients across samples from correlating the scaled activity of "(Non)mutational Epigenetic Reprogramming" with that of each pathway from which the genes were collected to represent the Hallmark signature. The annotation layers refer to the tumor types and the number of genes composing each pathway. **C)** Barplots for the number of pathways (left) and genes (right) that are finally obtained from the PathwayCommons database to be assigned for each Cancer Hallmark. **D)** Barplot for the number of associated hallmarks per gene. **E)** Boxplot for the percentage of overlap between each hallmark-associated gene list and the captured, post-filtered genes in each sample. **F)** Barplots for the 5 replicates of the 25% (top) and 50% (bottom) excluded percentage of genes from each pathway shown at the level of each hallmark-associated gene list. **G)** Heatmaps for the Pearson correlation coefficients from either gene exclusion (top) or gene addition (bottom) control experiments of the correlation between the quantified module score of the original signature of each hallmark and the average of that of the replicates for each corresponding hallmark within each control experiment. The corresponding standard deviations are shown in a heatmap to the right of the correlation coefficients' heatmaps. **H)** Boxplots for Moran's I values of the quantified module score of the original signature of each hallmark compared to those of randomly generated signatures of various gene sizes. ns: $p > 0.05$, *: $p <= 0.05$, **: $p <= 0.01$, ***: $p <= 0.001$, ****: $p <= 0.0001$

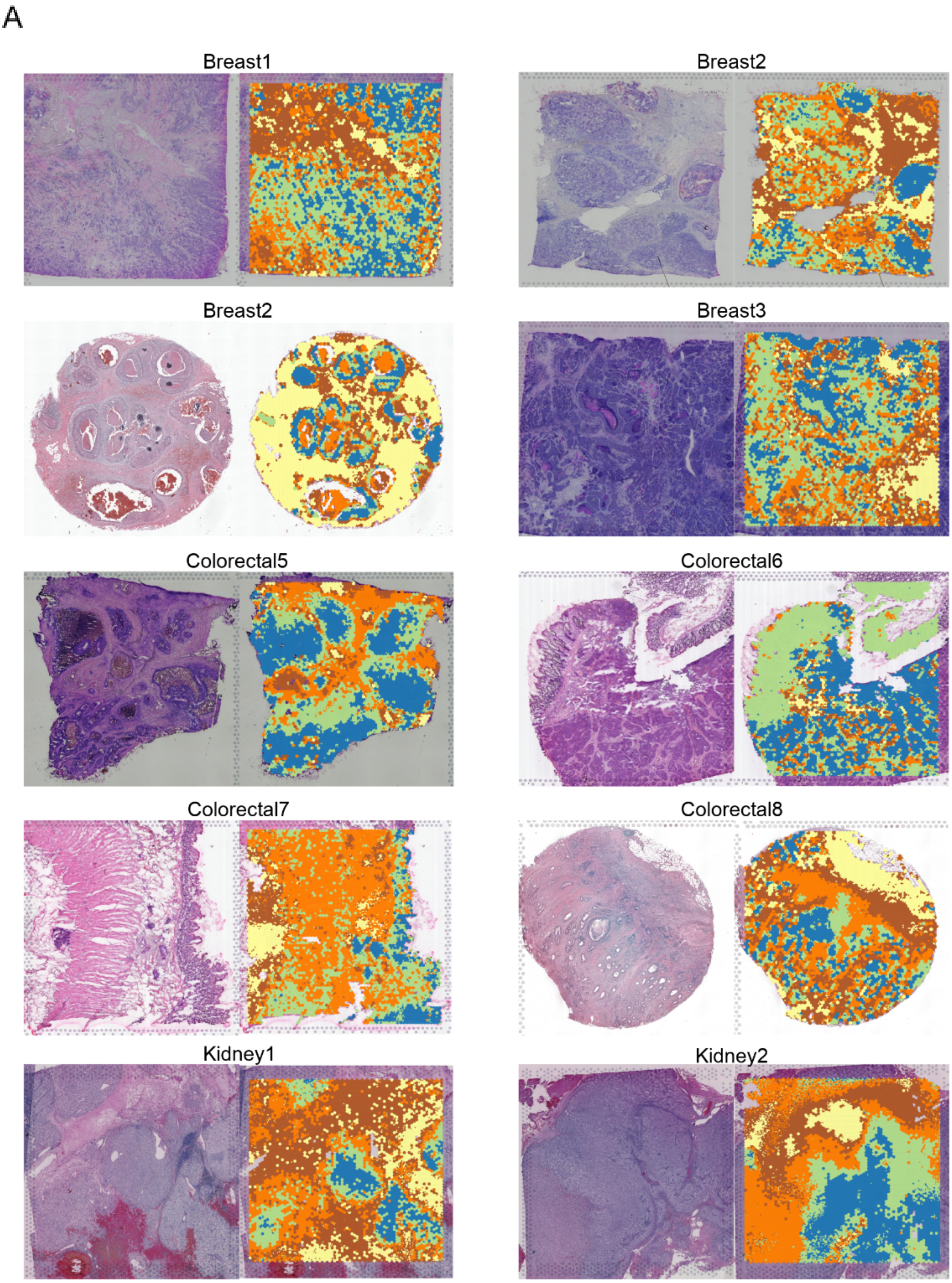
A
Breast1
Breast2
Breast2
Breast3
Colorectal5
Colorectal6
Colorectal7
Colorectal8
Kidney1
Kidney2

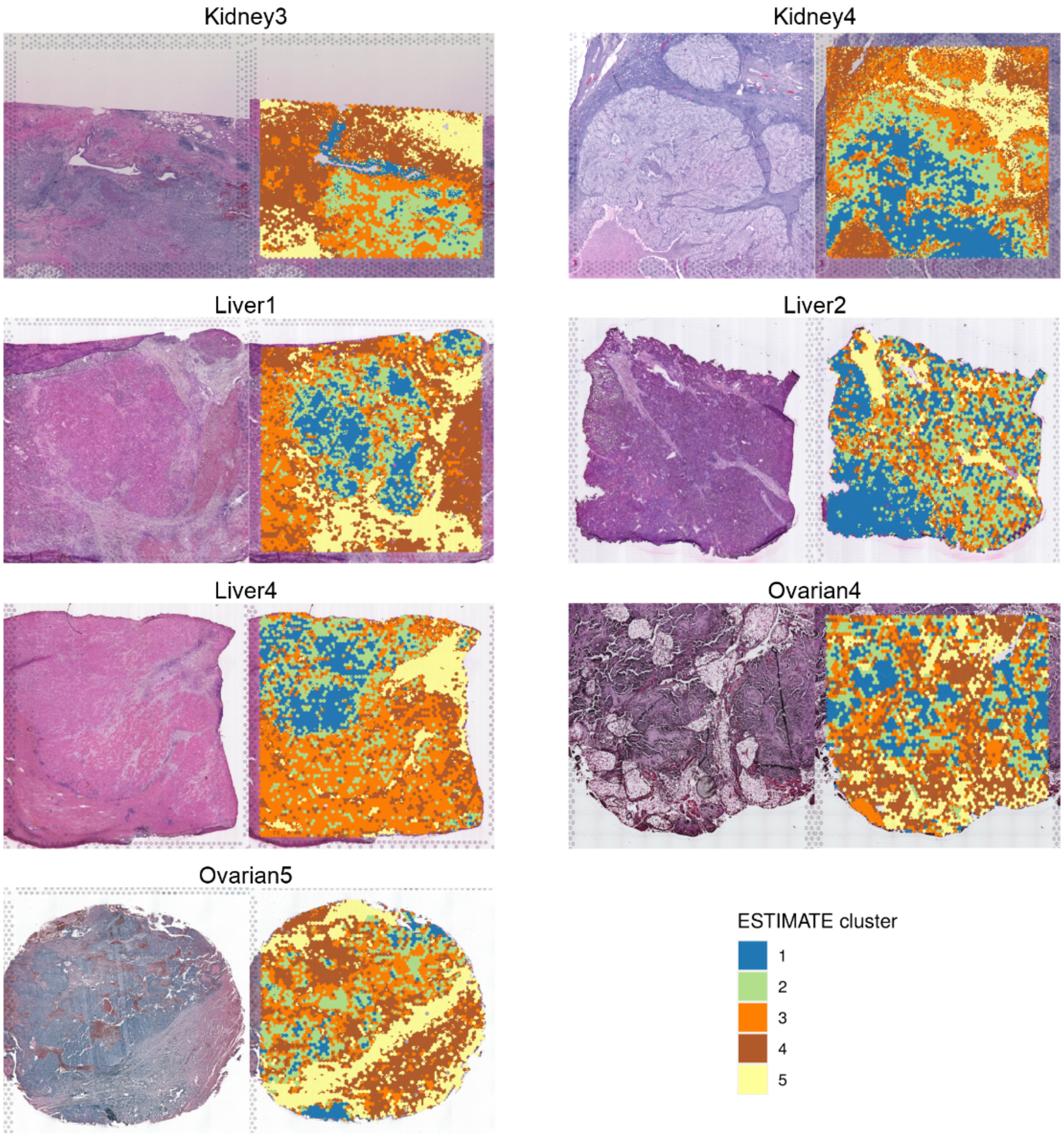

Kidney3
Kidney4
Liver1
Liver2
Liver4
Ovarian4
Ovarian5
ESTIMATE cluster
1
2
3
4
5

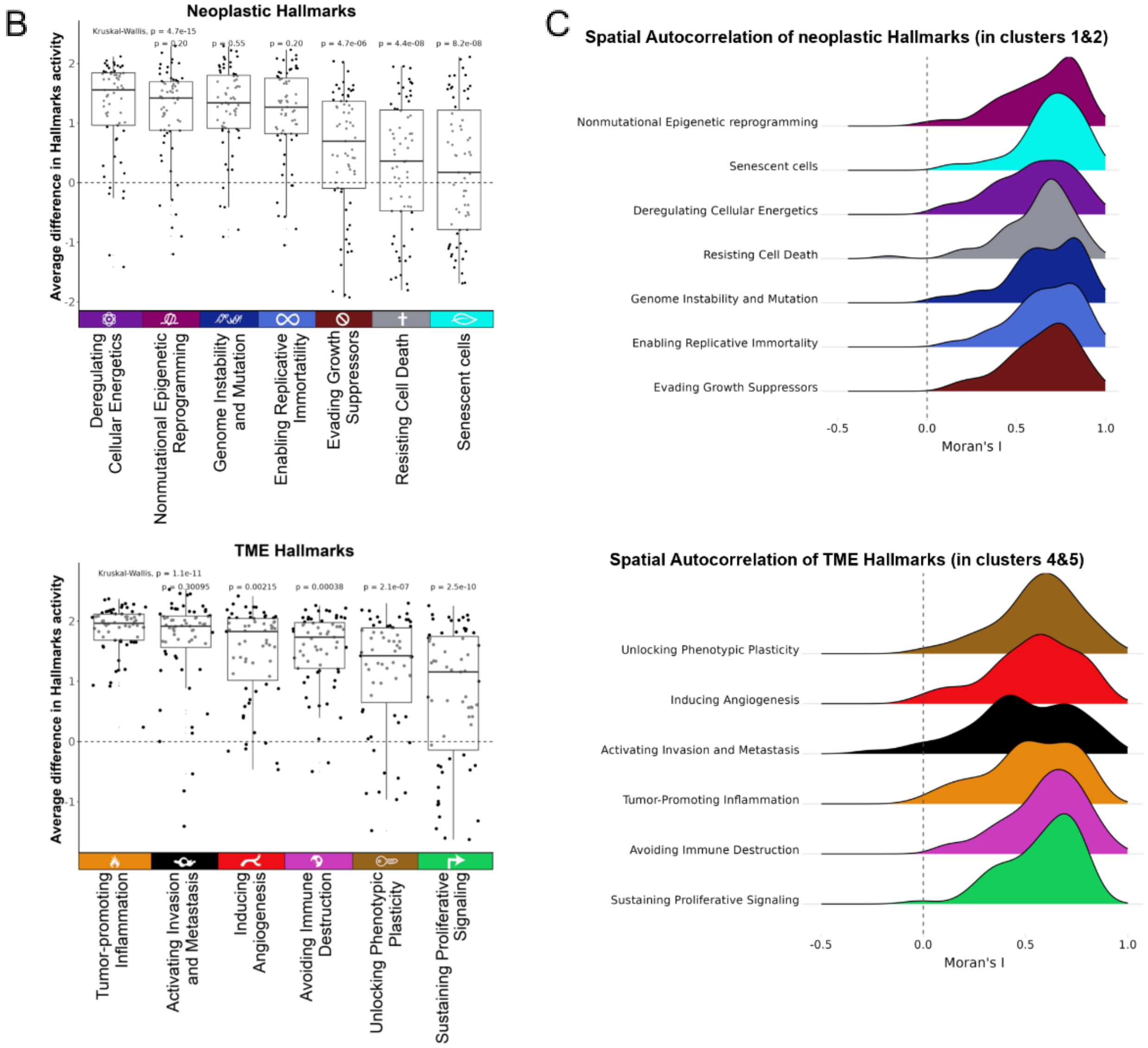

B
Neoplastic Hallmarks
Kruskal-Wallis, p = 4.7e-15
p = 0.20
p = 0.55
p = 0.20
p = 4.7e-06
p = 4.4e-08
p = 8.2e-08
Average difference in Hallmarks activity
Deregulating Cellular Energetics
Nonmutational Epigenetic Reprogramming
Genome Instability and Mutation
Enabling Replicative Immortatility
Evading Growth Suppressors
Resisting Cell Death
Senescent cells
TME Hallmarks
Kruskal-Wallis, p = 1.1e-11
p = 0.30095
p = 0.00215
p = 0.00038
p = 2.1e-07
p = 2.5e-10
Average difference in Hallmarks activity
Tumor-promoting Inflammation
Activating Invasion and Metastasis
Inducing Angiogenesis
Avoiding Immune Destruction
Unlocking Phenotypic Plasticity
Sustaining Proliferative Signaling
C
Spatial Autocorrelation of neoplastic Hallmarks (in clusters 1&2)
Nonmutational Epigenetic reprogramming
Senescent cells
Deregulating Cellular Energetics
Resisting Cell Death
Genome Instability and Mutation
Enabling Replicative Immortality
Evading Growth Suppressors
Moran's I
Spatial Autocorrelation of TME Hallmarks (in clusters 4&5)
Unlocking Phenotypic Plasticity
Inducing Angiogenesis
Activating Invasion and Metastasis
Tumor-Promoting Inflammation
Avoiding Immune Destruction
Sustaining Proliferative Signaling
Moran's I

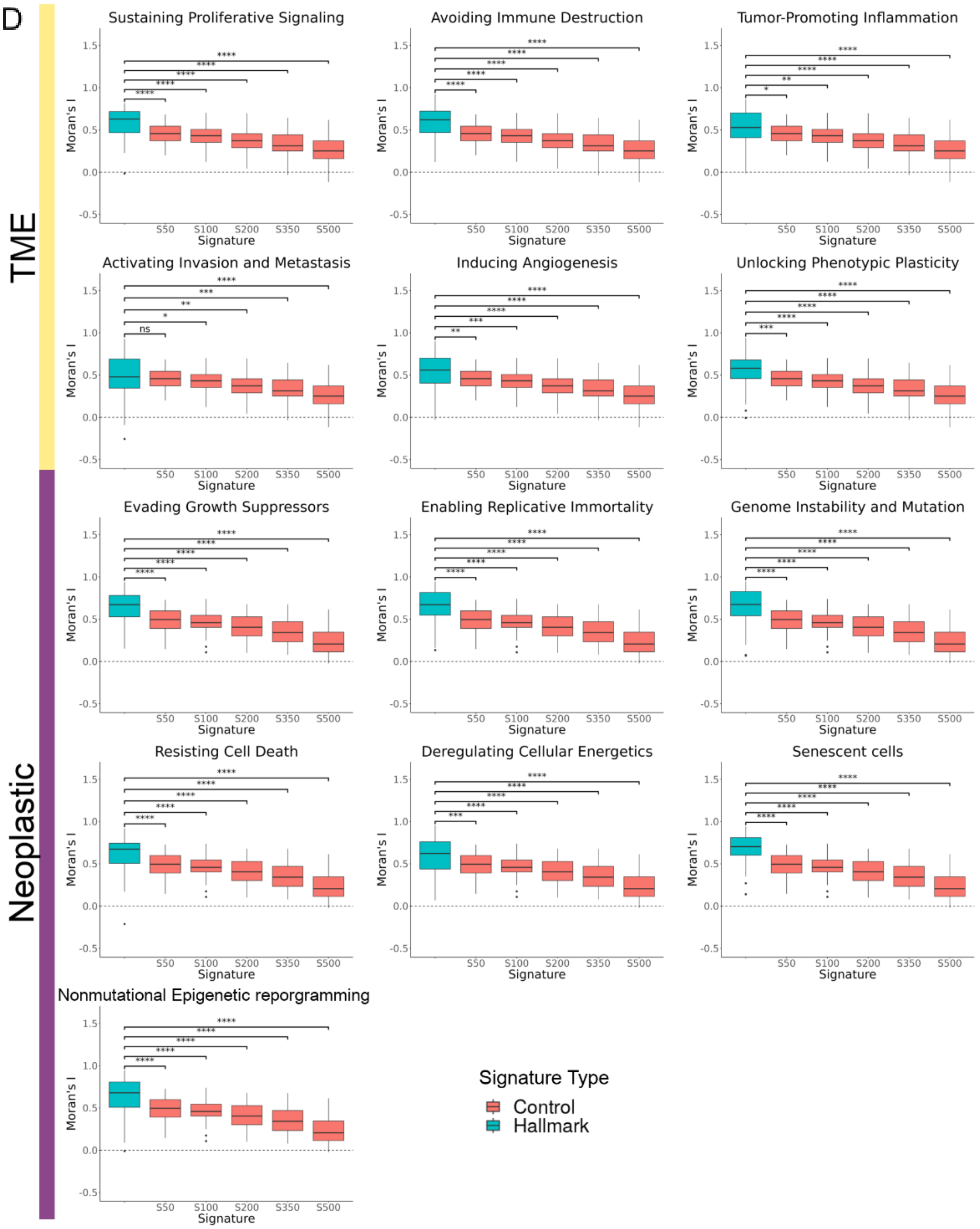


**Figure S2: ESTIMATE clusters, Hallmark differences between tumor compartments, and their corresponding spatial autocorrelation along with their controls. A)** H&E images (left) and the ESTIMATE clusters (right) overlaid on the histology image of 17 samples from 5 cancer types that were used by expert human pathologists to validate the cancer cell purity of the ESTIMATE scores. **B)** The average difference of hallmark activities between neoplastic (clusters 1&2) and TME (clusters 4&5) compartments. Positive values indicate higher neoplastic hallmark activities in the neoplastic compartment (top) or higher

TME Hallmarks in the TME compartment (bottom). **C)** Ridge plots of Moran's I computed for neoplastic hallmarks within the neoplastic compartment (top) and for TME hallmarks within TME compartment (bottom). **D)** Boxplots for Moran's I values of the quantified module score of the original signature of the TME hallmarks in the TME compartment (top) and neoplastic hallmarks in the neoplastic compartment (bottom) compared to those of randomly generated signatures of various gene sizes. ns: p > 0.05, *: p <= 0.05, **: p <= 0.01, ***: p <= 0.001, ****: p <= 0.0001

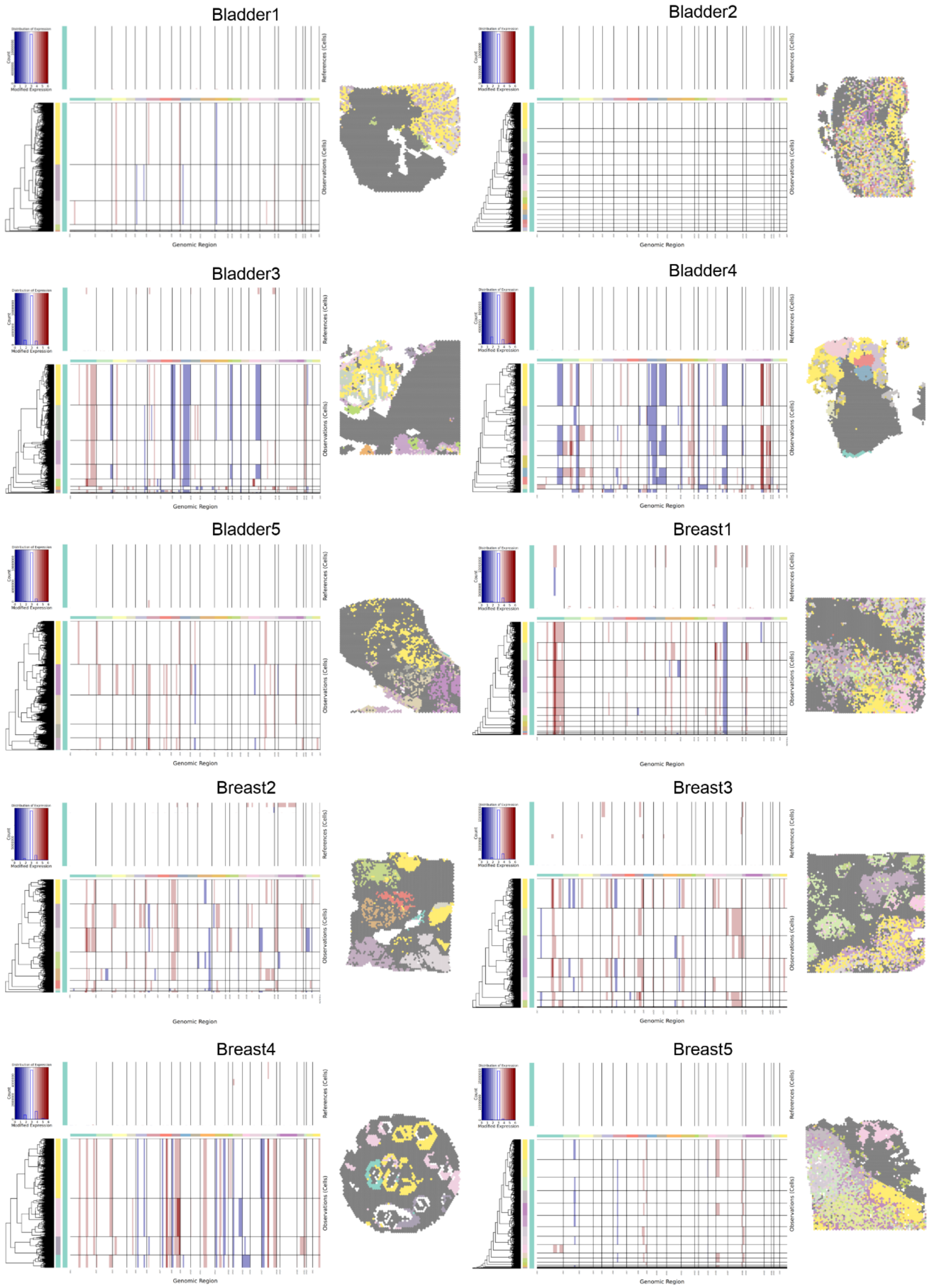
Bladder1
Bladder2
Bladder3
Bladder4
Bladder5
Breast1
Breast2
Breast3
Breast4
Breast5
Modified Expression
Count
References (Cells)
Observations (Cells)
Genomic Region

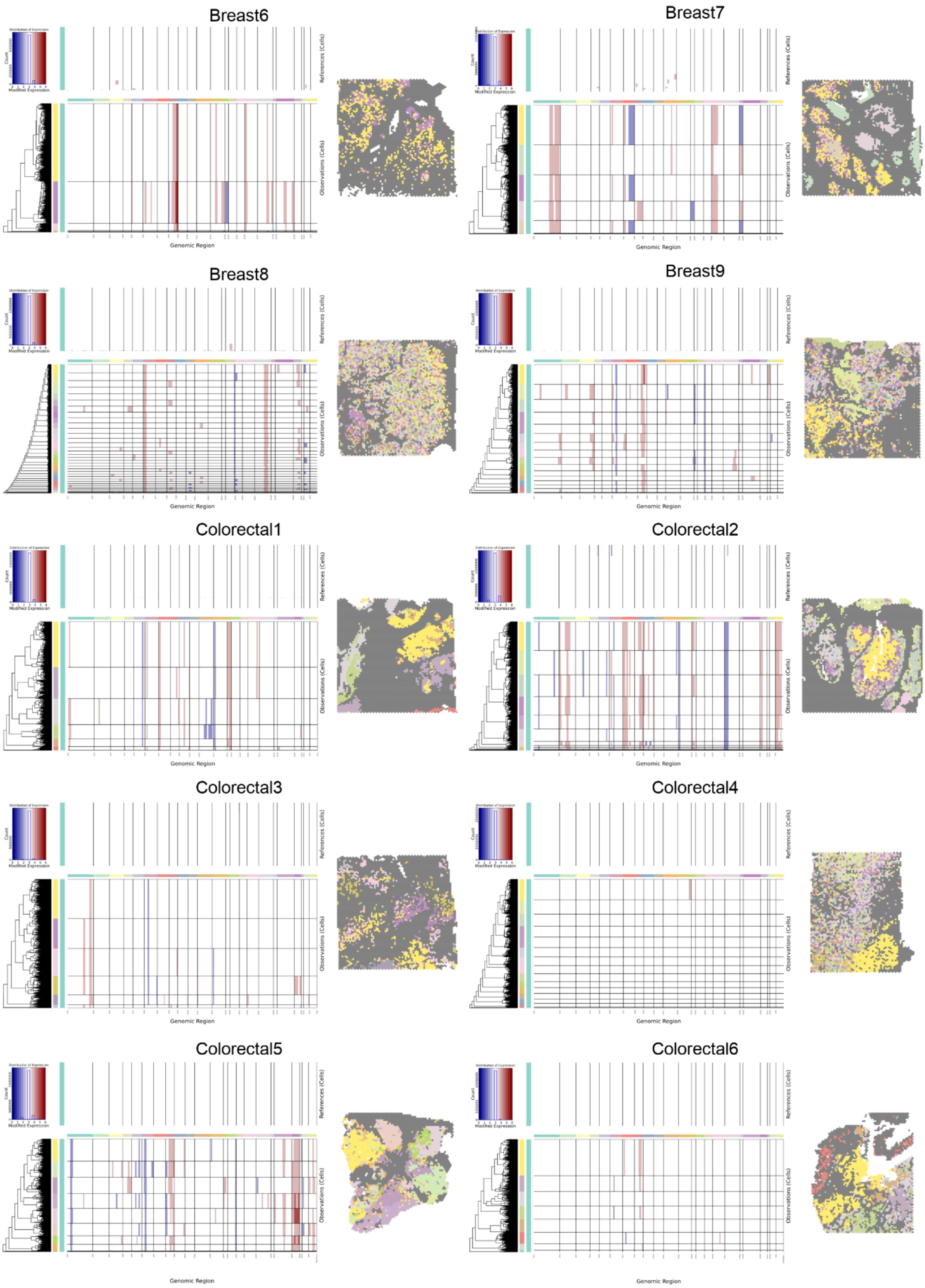
Breast6
Breast7
Breast8
Breast9
Colorectal1
Colorectal2
Colorectal3
Colorectal4
Colorectal5
Colorectal6
Modified Expression
Count
References (Cells)
Observations (Cells)
Genomic Region

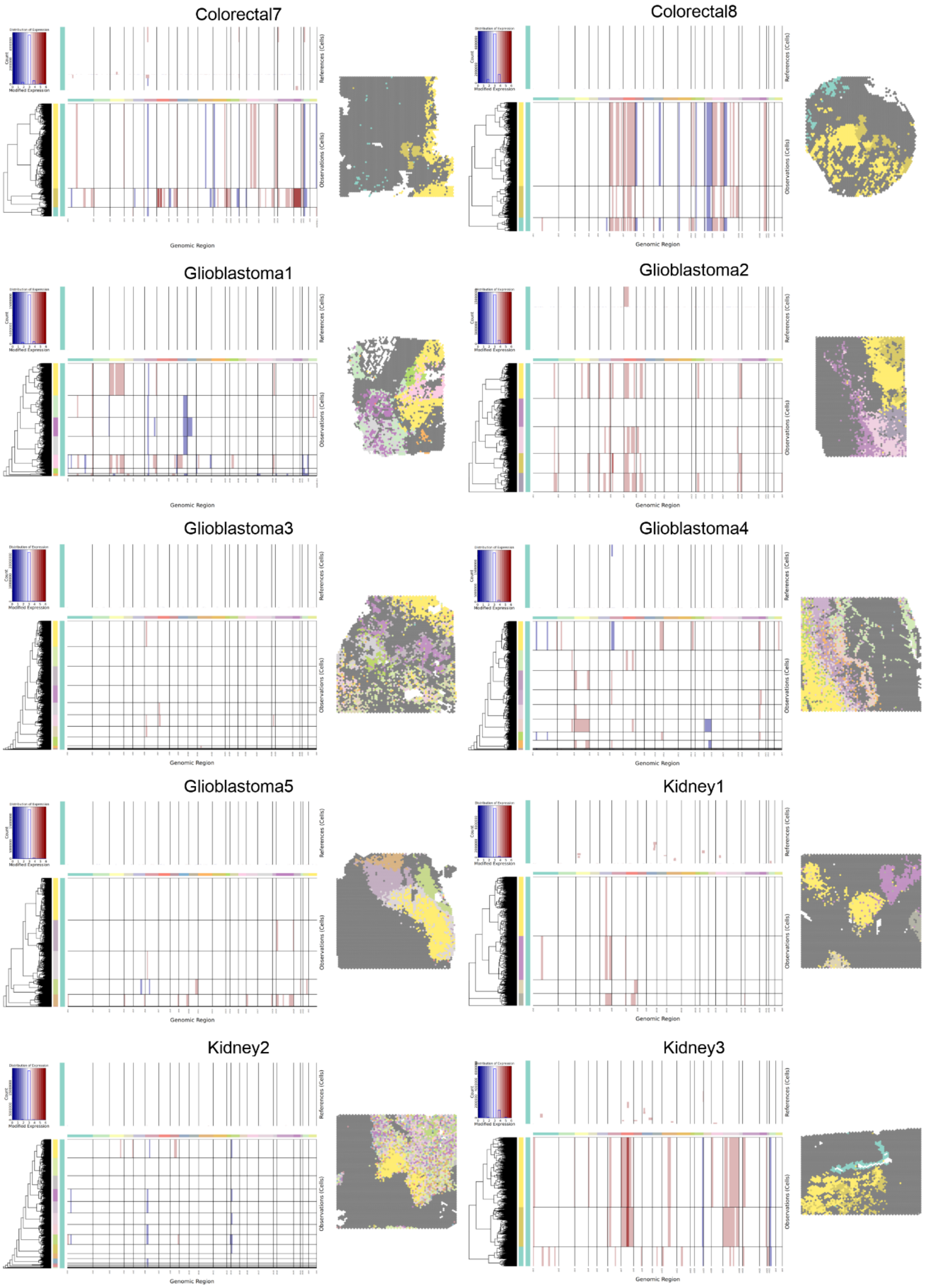
Colorectal7
Colorectal8
Glioblastoma1
Glioblastoma2
Glioblastoma3
Glioblastoma4
Glioblastoma5
Kidney1
Kidney2
Kidney3
References (Cells)
Observations (Cells)
Genomic Region
Count
Modified Expression

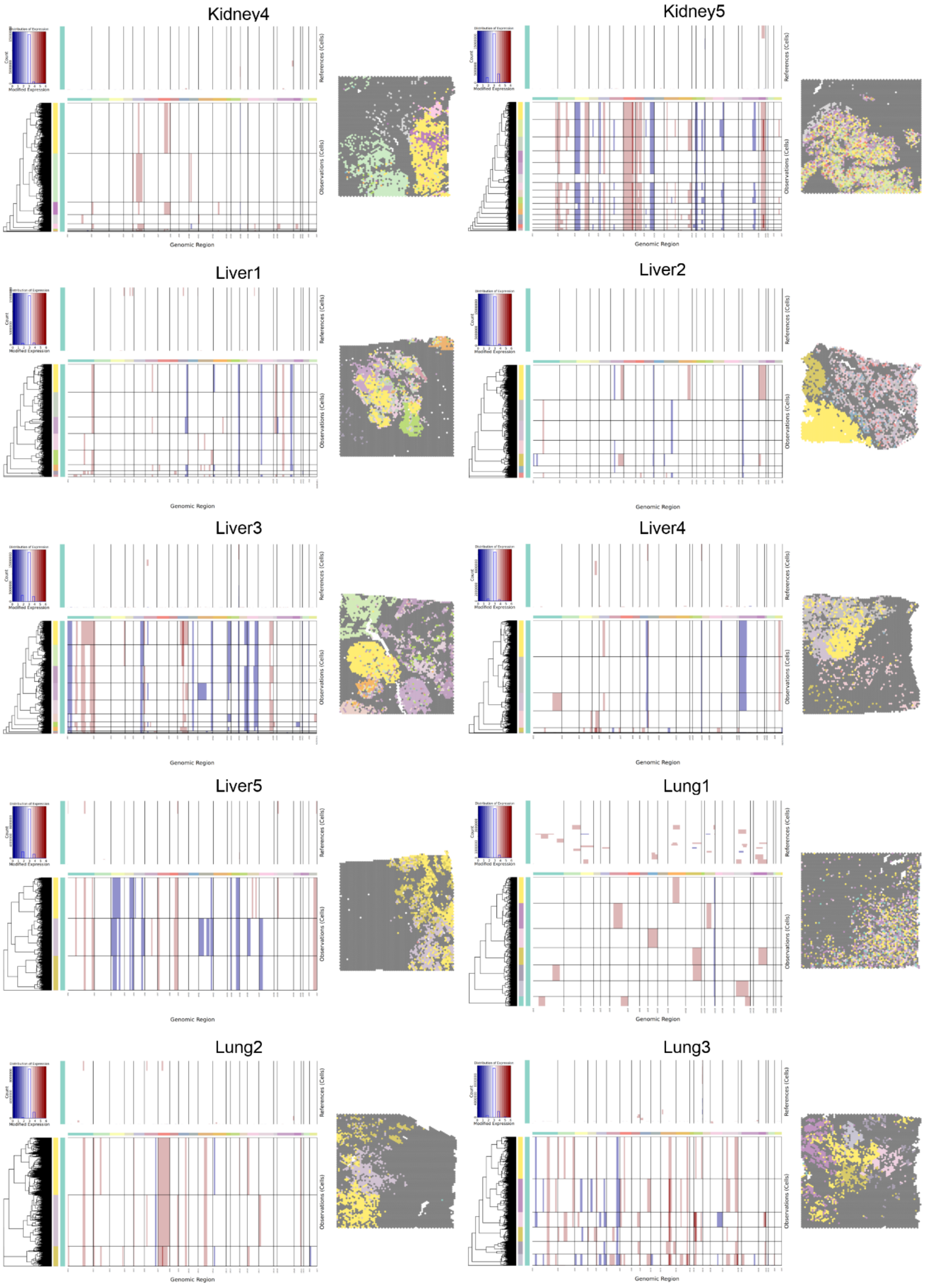
Kidney4
Kidney5
Liver1
Liver2
Liver3
Liver4
Liver5
Lung1
Lung2
Lung3
Modified Expression
References (Cells)
Observations (Cells)
Genomic Region

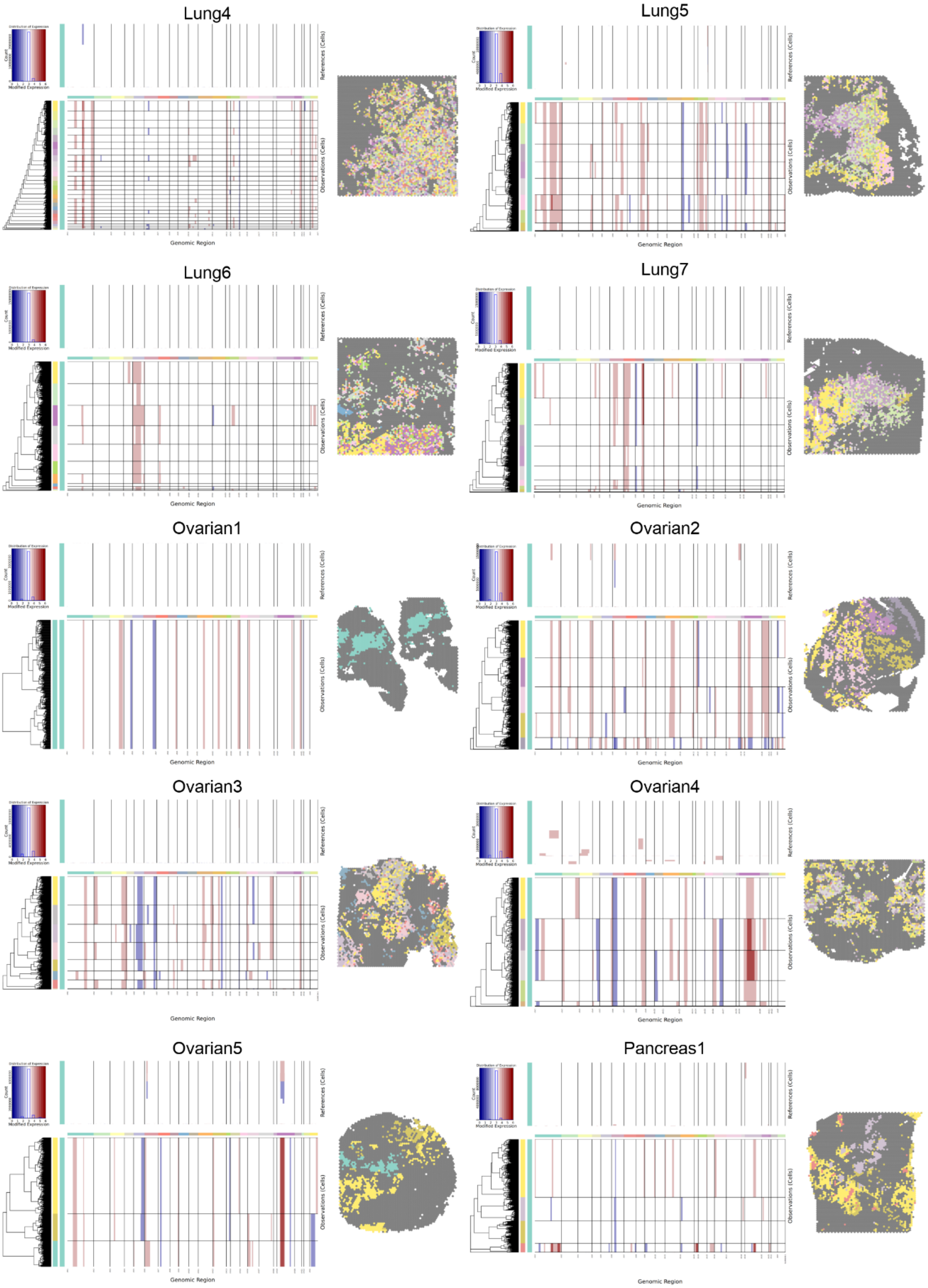
Lung4
Lung5
Lung6
Lung7
Ovarian1
Ovarian2
Ovarian3
Ovarian4
Ovarian5
Pancreas1
Modified Expression
References (Cells)
Observations (Cells)
Genomic Region

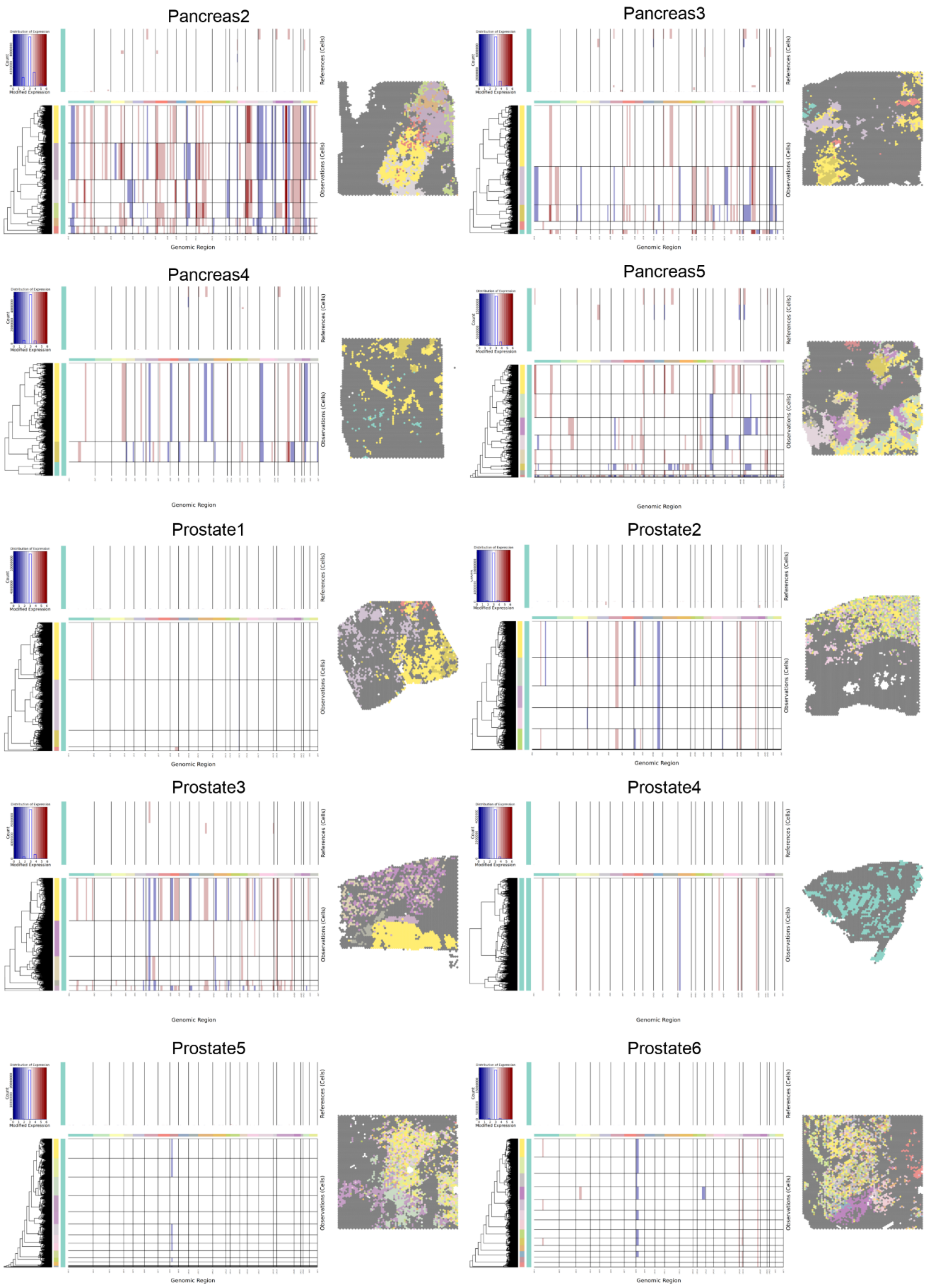
Pancreas2
Pancreas3
Pancreas4
Pancreas5
Prostate1
Prostate2
Prostate3
Prostate4
Prostate5
Prostate6
Distribution of Expression
Count
Modified Expression
References (Cells)
Observations (Cells)
Genomic Region

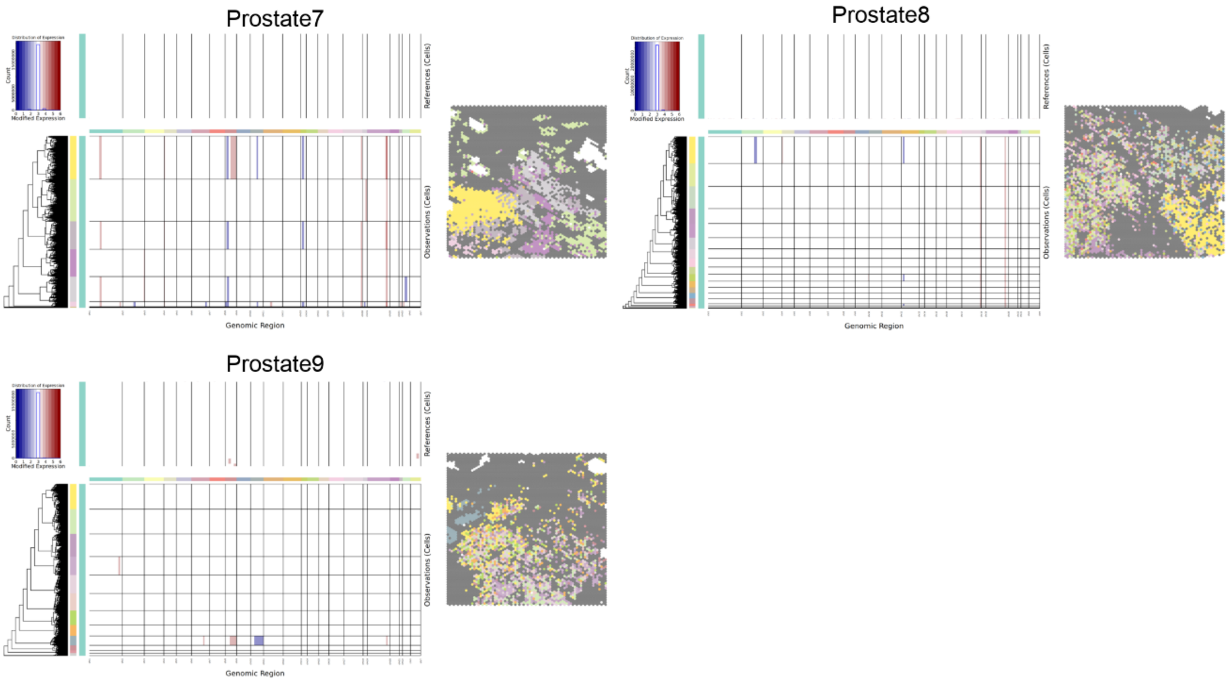


**Figure S3: Predicted clonal architecture reported by inferCNV.** Heatmap of clusters of chromosomal gain and loss from the HMM model generated by inferCNV that are used to predict a number of clones. The predicted clones are projected on the ST data shown to the right of the heatmap for each of the 63 samples.

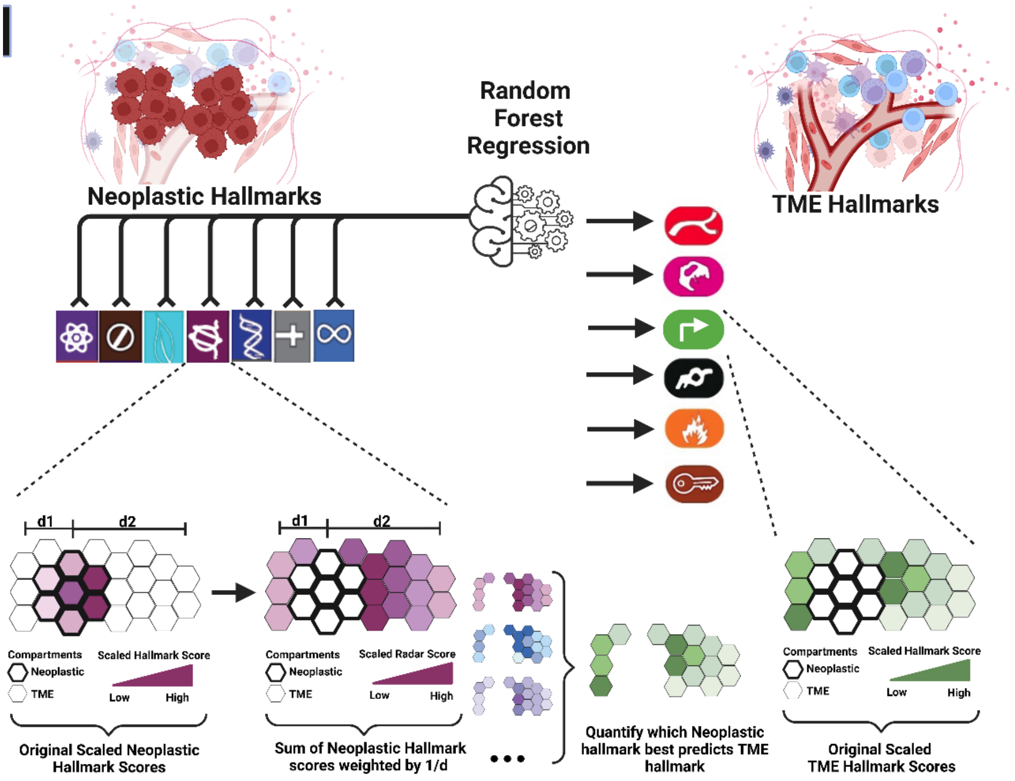


**Figure S4: Schematic diagram illustrating the process of using random forest to predict the spatial landscape of each TME hallmark.** Illustration of using Random Forest regression to predict TME Hallmarks within the TME compartment using spatial distribution of Hallmarks from the neoplastic compartment. This is accomplished by means of transforming the activities of neoplastic hallmarks into "radar" scores from their associated compartment into the TME compartment where these radars will act as the predictors in the random forest model.

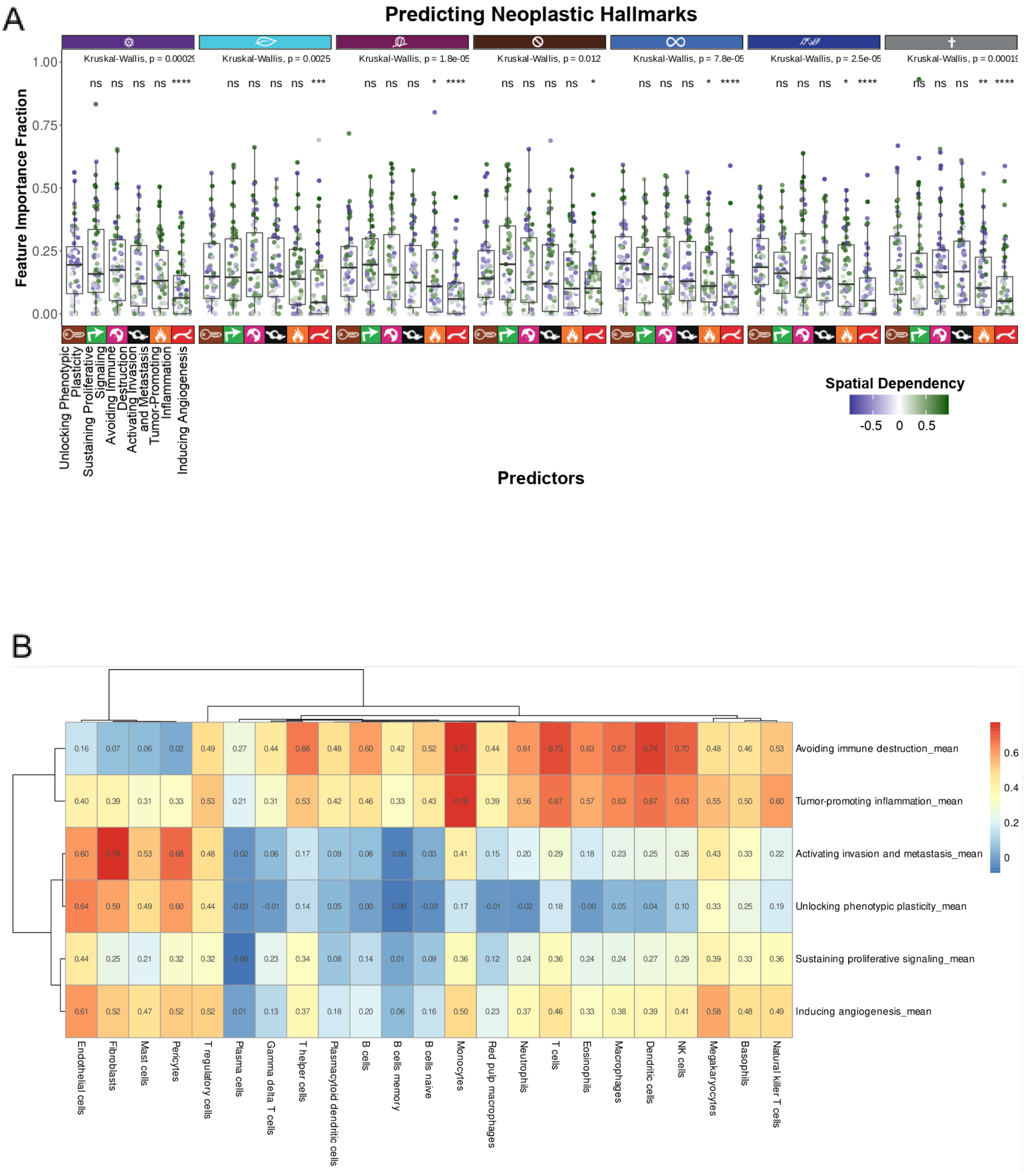
A
Predicting Neoplastic Hallmarks
Feature Importance Fraction
1.00
0.75
0.50
0.25
0.00
Kruskal-Wallis, p = 0.00029
Kruskal-Wallis, p = 0.0025
Kruskal-Wallis, p = 1.8e-05
Kruskal-Wallis, p = 0.012
Kruskal-Wallis, p = 7.8e-05
Kruskal-Wallis, p = 2.5e-05
Kruskal-Wallis, p = 0.00019
Unlocking Phenotypic Plasticity
Sustaining Proliferative Signaling
Avoiding Immune Destruction
Activating Invasion and Metastasis
Tumor-Promoting Inflammation
Inducing Angiogenesis
Predictors
Spatial Dependency
-0.5 0 0.5
B
Avoiding immune destruction_mean
Tumor-promoting inflammation_mean
Activating invasion and metastasis_mean
Unlocking phenotypic plasticity_mean
Sustaining proliferative signaling_mean
Inducing angiogenesis_mean
Endothelial cells
Fibroblasts
Mast cells
Pericytes
T regulatory cells
Plasma cells
Gamma delta T cells
T helper cells
Plasmacytoid dendritic cells
B cells
B cells memory
B cells naive
Monocytes
Red pulp macrophages
Neutrophils
T cells
Eosinophils
Macrophages
Dendritic cells
NK cells
Megakaryocytes
Basophils
Natural killer T cells
0.6
0.4
0.2
0

C

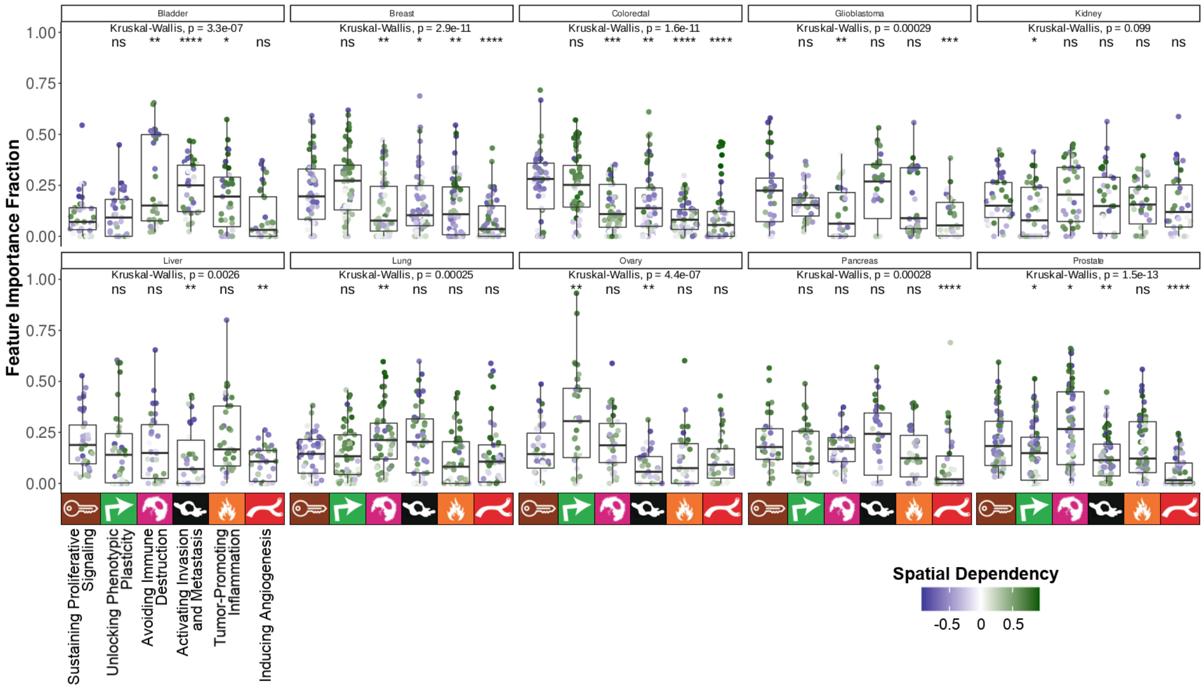


**Figure S5: The relative importance of TME Hallmarks as predictors of neoplastic Hallmarks faceted by target and tumor type. A)** Boxplot showing the feature importance fraction of TME hallmarks (their radars) when predicting each target neoplastic hallmark in each of the 63 samples. Each dot of these plots is colored by the spatial dependency between the predictor and target pairs. **B)** Heatmap depicts the average Pearson correlation coefficients across samples from correlating the individual activities of each of the 6 TME Hallmarks with the quantified scores of over 20 different cell types within the TME compartment. **C)** the same information presented in A is presented here when facetting by cancer type. ns: p > 0.05, *: p <= 0.05, **: p <= 0.01, ***: p <= 0.001, ****: p <= 0.0001

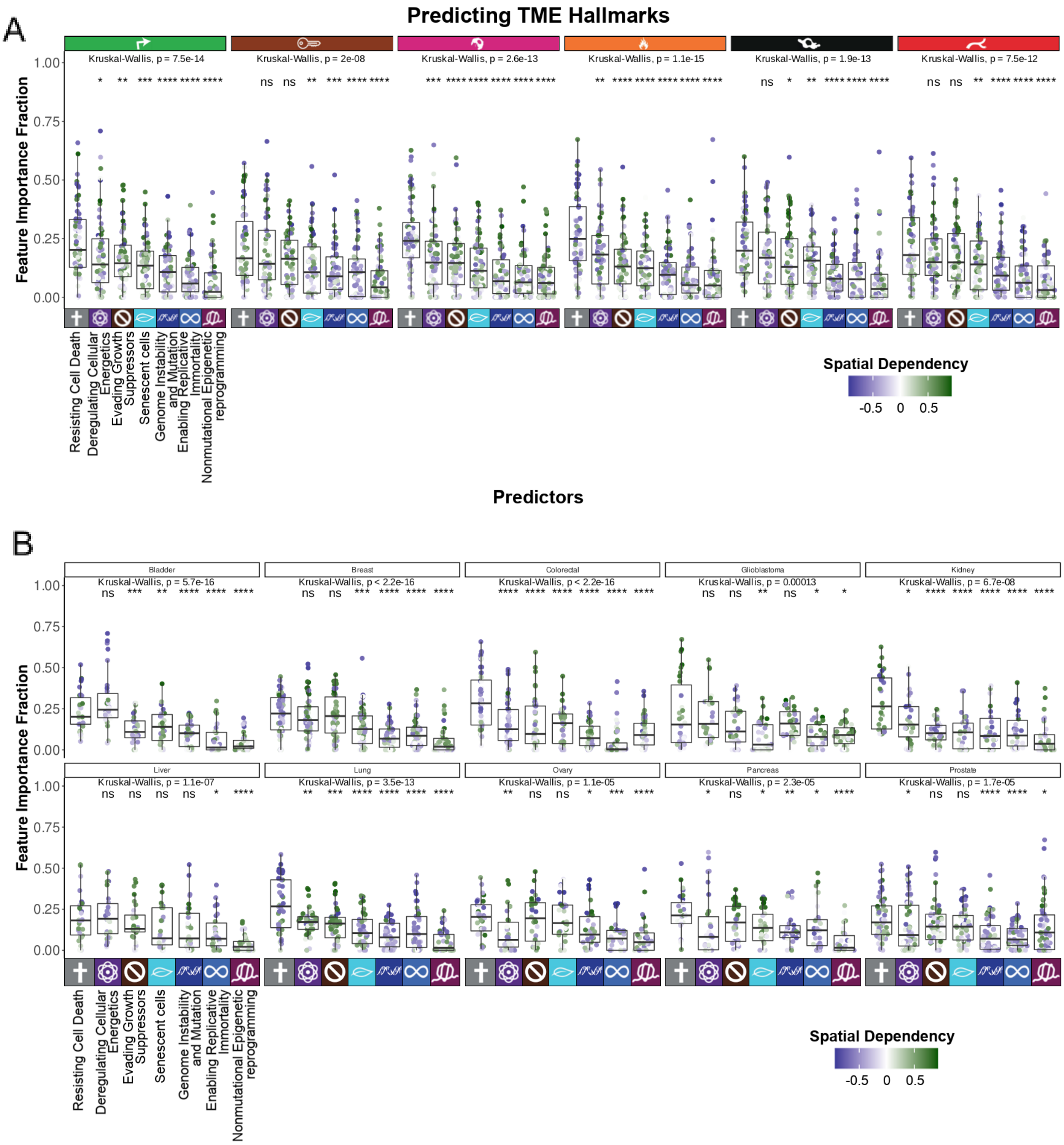


**Figure S6: The relative importance of neoplastic Hallmarks as predictors of TME Hallmarks faceted by target and tumor type. A)** Boxplot showing the feature importance fraction of neoplastic hallmarks (their radars) when predicting each target TME hallmark in each of the 63 samples. Each dot of these plots is colored by the spatial dependency between the predictor and target pairs. **B)** The same information presented in A is presented here when facetting by cancer type. ns: p > 0.05, *: p <= 0.05, **: p <= 0.01, ***: p <= 0.001, ****: p <= 0.0001
